# Behavioral pattern separation and cognitive flexibility are enhanced in a mouse model of increased lateral entorhinal cortex-dentate gyrus circuit activity

**DOI:** 10.1101/2023.01.26.525756

**Authors:** Sanghee Yun, Ivan Soler, Fionya Tran, Harley A. Haas, Raymon Shi, Grace L. Bancroft, Maiko Suarez, Chris R. de Santis, Ryan P. Reynolds, Amelia J. Eisch

## Abstract

Behavioral pattern separation and cognitive flexibility are essential cognitive abilities which are disrupted in many brain disorders. Better understanding of the neural circuitry involved in these abilities will open paths to treatment. In humans and mice, discrimination and adaptation rely on integrity of the hippocampal dentate gyrus (DG) which both receive glutamatergic input from the entorhinal cortex (EC), including the lateral EC (LEC). Inducible increase of EC-DG circuit activity improves simple hippocampal-dependent associative learning and increases DG neurogenesis. Here we asked if the activity of LEC fan cells that directly project to the DG (LEC➔DG neurons) regulates behavioral pattern separation or cognitive flexibility. C57BL6/J male mice received bilateral LEC infusions of a virus expressing shRNA TRIP8b, an auxiliary protein of an HCN channel or a control virus (SCR shRNA); this approach increases the activity of LEC➔DG neurons. Four weeks later, mice underwent testing for behavioral pattern separation and reversal learning (touchscreen-based Location Discrimination Reversal [LDR] task) and innate fear of open spaces (elevated plus maze [EPM]) followed by counting of new DG neurons (doublecortin-immunoreactive cells [DCX+] cells). TRIP8b and SCR shRNA mice performed similarly in general touchscreen training and LDR training. However, in late LDR testing, TRIP8b shRNA mice reached the first reversal more quickly and had more accurate discrimination vs. SCR shRNA mice, specifically when pattern separation was challenging (lit squares close together or “small separation”). Also, TRIP8b shRNA mice achieved more reversals in late LDR testing vs. SCR shRNA mice. Supporting a specific influence on cognitive behavior, SCR shRNA and TRIP8b shRNA mice did not differ in total distance traveled or in time spent in the closed arms of the EPM. Supporting an inducible increase in LEC-DG activity, DG neurogenesis was increased. These data indicate TRIP8b shRNA mice had better pattern separation and reversal learning and more neurogenesis vs. SCR shRNA mice. This work advances fundamental and translational neuroscience knowledge relevant to two cognitive functions critical for adaptation and survival — behavioral pattern separation and cognitive flexibility — and suggests the activity of LEC➔DG neurons merits exploration as a therapeutic target to normalize dysfunctional DG behavioral output.

## INTRODUCTION

Successful adaptation and survival demand the ability to discriminate stimuli (learn which stimulus is associated with a reward) and to be flexible (change behavior when reward conditions change)(Yassa and Stark, 2011; Dajani and Uddin, 2015; Prado et al., 2017; Cayco-Gajic and Silver, 2019). An example of stimuli discrimination is behavioral pattern separation, where a subject learns over time which stimulus (S) is associated with a meaningful outcome, such as reward delivery (S+). When stimuli or episodes are very similar, and thus cognitive load is high (Yassa et al., 2011; Bekinschtein et al., 2013; Kent et al., 2015a, 2015b; Kassab and Alexandre, 2018), this discrimination learning can be considered a behavioral readout of *pattern separation*, a process by which the brain avoids confusion between similar memories (Rolls and Kesner, 2006; Schmidt et al., 2012; Kesner and Rolls, 2015; Anacker and Hen, 2017; Severa et al., 2017; Cayco-Gajic and Silver, 2019). A key component of flexible learning is reversal learning, where a subject adapts its behavior when outcome conditions change (as when S+ becomes S-). Reversal learning is a behavioral readout of *cognitive flexibility* which, together with strategy shifting, allows successful adaptation to a changing world (Bissonette and Powell, 2012; Izquierdo et al., 2017). Many studies suggest behavioral pattern separation and cognitive flexibility engage and rely on the integrity of hippocampal circuitry (Leutgeb et al., 2007; Clelland et al., 2009; Yassa and Stark, 2011; Burghardt et al., 2012; Swan et al., 2014; Anacker and Hen, 2017). Far less is known about the role of a brain region that provides direct input to the DG: the entorhinal cortex (EC).

The upstream EC is considered a functional gatekeeper for the downstream hippocampus (Fernandez and Tendolkar, 2006; Basu et al., 2016; Hansen et al., 2018), feeding multi-sensory and associative information into the DG and other hippocampal regions (Bekinschtein et al., 2013; Morrissey and Takehara-Nishiuchi, 2014; Reagh and Yassa, 2014; Kitamura et al., 2015; Save and Sargolini, 2017; Reagh et al., 2018; Morales et al., 2021; Marks et al., 2022). In addition to the EC being involved in spatial navigation and episodic memory, one study suggests the EC — and particularly the lateral EC (LEC) — is critical for behavioral pattern separation (Vivar et al., 2012). Using the touchscreen-based Location Discrimination Reversal (LDR) task, Vivar and colleagues reported that male mice who received LEC excitotoxic lesions took more trials to reach task criterion compared to mice that received control infusions. A more recent study used cell ablation to test a role for LEC activity in making associations crucial for behavioral pattern separation (Vandrey et al., 2020). Specifically, cre-mediated ablation of LEC layer IIa (LECIIa) fan cells in male mice impaired complex associative episodic learning (e.g. object-place-context) but not simpler associative learning (Vandrey et al., 2020). LECIIa fan cells are notable as they comprise the only projection from the EC to the DG; they make excitatory synapses in the DG molecular layer (Mol) on the processes of glutamatergic DG granule cells (GCs), adult-born GCs, and mossy cells, and GABAergic interneurons (Witter, 2007; Witter et al., 2017; Vandrey et al., 2020; Traub and Whittington, 2022). All of these DG cell types have been implicated in behavioral pattern separation (Leutgeb et al., 2007; Myers and Scharfman, 2009; Jinde et al., 2012; Schmidt et al., 2012; GoodSmith et al., 2017; Nakazawa, 2017; Morales et al., 2021), and some of them — including adult-born DG GCs — have also been implicated in cognitive flexibility (Burghardt et al., 2012; Swan et al., 2014; Yagi and Galea, 2019; Wingert and Sorg, 2021). This circuit positioning of LECIIa fan cells suggests they are a “load-sensitive” component of the EC-DG circuit in behavioral pattern separation and cognitive flexibility, but this has not yet been tested.

We hypothesized stimulation of LECIIa fan cells, and thus stimulation of the LEC➔DG projection, would improve behavioral pattern separation and cognitive flexibility. To test this, we modified an approach previously used to stimulate both LECIIa fan cells and medial EC (MEC) layer IIa stellate cells (Yun et al., 2018a). Specifically, we used viral-mediated knock down via shRNA of tetratricopeptide repeat-containing Rab8b-interacting protein (TRIP8b), a brain-specific auxiliary subunit of the Hyperpolarization-Activated Cyclic Nucleotide-Gated Channel (HCN) (Yun et al., 2018a). Relative to control shRNA mice, mice that had EC TRIP8b shRNA in ECIIa fan/stellate cells had several indices of increased EC activity: more firing in EC cells that projected to the DG and hippocampus, higher levels of DG activity-dependent processes (e.g. DG neurogenesis, dendritic arborization), and improved hippocampal-dependent contextual memory. Here we narrowed our focus to the LEC to fill the knowledge gap on whether behavioral pattern separation and cognitive flexibility are influenced by the activity of LECIIa fan cells that project to the DG (LEC➔DG neurons). We report that relative to control mice, mice with TRIP8b shRNA in LECIIa fan cells had improved behavioral pattern separation and cognitive flexibility as well as more DG neurogenesis. Our findings clarify the circuitry engaged in these cognitive abilities, implicating the activity of the “upstream” LEC for the first time in behavioral pattern separation and cognitive flexibility.

## METHODS AND MATERIALS

### Animals and Ethics Statement

Experiments were approved by the Institutional Animal Care and Use Committee at the Children’s Hospital of Philadelphia (CHOP) and performed in compliance with the National Institutes of Health Guide for the Care and Use of Laboratory Animals. Mice were group-housed by treatment in an AAALAC-accredited, specific-pathogen free conventional vivarium at the CHOP. Six-week-old C57BL/6J male mice were purchased from Jackson Laboratory (stock number: 000664) and housed in a CHOP vivarium for at least one week prior to the start of the study. Room environments were maintained according to Guide standards (20-23°C and 30-70% humidity). Home cages consisted of individually ventilated polycarbonate microisolator cages (Lab Products Inc., Enviro-Gard™ III, Seaford, DE) with HEPA filtered air, corncob (Bed-o’ Cobs® ¼”) bedding, with provision of one nestlet (Ancare) and a red plastic hut (Bio-Serv, #K3583 Safe Harbor). Each cage of 4 mice/cage was randomly-assigned to the scrambled shRNA (SCR) or TRIP8b shRNA group. Individual mice were identified using the ear punch system. Mice were kept on a 12-hour (h) light/dark cycle (lights on at 06:15) with *ad libitum* access to chow (Lab Diets 5015 #0001328) and water. After starting touchscreen experiments, mice were given access to chow from 13:30 to 17:30 (4h/day [d]) Monday through Thursday and *ad libitum* access from Friday 13:30 to Sunday 17:30. Mouse weight was recorded at surgery and weekly thereafter to ensure mice remain above 85% of their initial weight. The data from n=2 SCR shRNA mice were excluded from this experiment, as detailed in the <u>Rigor, Additional ARRIVE 2.0 Details, and Statistical Analysis</u> section below.

### AAV Vectors

shRNAs were designed to target the TRIP8b C-terminus region (exon 11) as previously published (Lewis et al., 2009). The following oligonucleotides were used with overhanging ends identical to those created by Sap I and Xba I restriction enzymes: For TRIP8b shRNA, 5’-

TTTGAGCATTTGAAGAAGGCTTAATTCAAGAGATTAAGCCTT

CTTCAAAT GCTATTTTT-3’; SCR shRNA, 5’-TTTGTTCTCCGAACGT

GTCACGTTTCAAGAGAACGTGACACGTTCGGAGAATTTTT-3’. Hairpin oligonucleotides were phosphorylated by T4 polynucleotide kinase (New England Biolabs, Beverly, MA) followed by annealing at 100°C for 5min and cooling in the heat block for 3h. Each annealed oligonucleotide was ligated into the adeno-associated virus (AAV2) plasmid (pAAV-EGFP-shRNA; Stratagene, La Jolla, CA)(Hommel et al., 2003). Virus production was achieved from UPenn Vector core (Yun et al., 2018b).

### Stereotaxic Surgery

Mice were anesthetized with a mixture of ketamine (120 mg/kg) and xylazine (16 mg/kg) in saline (0.9% NaCl, i.p.). Bilateral stereotaxic injection of 0.4-0.5 µl of purified high titer AAV (TRIP8b shRNA or SCR shRNA) was directed into the LEC (from Bregma: A/P-3.7 mm, M/L+4.4, angle 4°; from Lambda: D/V-4.5) using 33 gauge Hamilton syringes (Hamilton, Reno, NV). Injection rate was 0.1 µl/min, with needles kept in place for an additional 5min to enable diffusion.

### Overview of Behavioral Testing

Mice with infusion of SCR shRNA or TRIP8b shRNA virus in the LEC (**Fig. 1A**) underwent touchscreen behavioral testing 1 week post-surgery. Touchscreen experiments were performed between 08:00 to 14:00 during weekdays (Soler et al., 2021). As is standard in most rodent touchscreen experiments, mice were food restricted during touchscreen experiments. Mouse chow was removed from each cage at 5 pm the day prior to training or testing. Each cage was given *ad libitum* access to chow for 3h (minimum) to 4h (maximum) immediately following daily touchscreen training/testing, and from completion of training/testing on Friday until Sunday 5 pm. Mice were weighed weekly to ensure weights >85% initial body weight. While weights below this threshold merited removal of the mouse from the study, zero mice reached this threshold (Mar et al., 2013; Oomen et al., 2013). Operant touchscreen platform procedures included general touchscreen training (with 2 × 6 window grid) and Location Discrimination Reversal [LDR] training and testing; LDR was selected for its ability to reflect an animal’s performance on both behavioral pattern separation and cognitive flexibility (Swan et al., 2014). After all mice completed LDR testing, mice received unrestricted food pellets for one week prior to behavior testing on the elevated plus maze (EPM) for innate fear/anxiety measurement. Subject number overview is provided in the Statistical Analysis section below, and the subject number for each group in each figure panel is provided in **Supplementary Table 1**.

**Figure 1.**
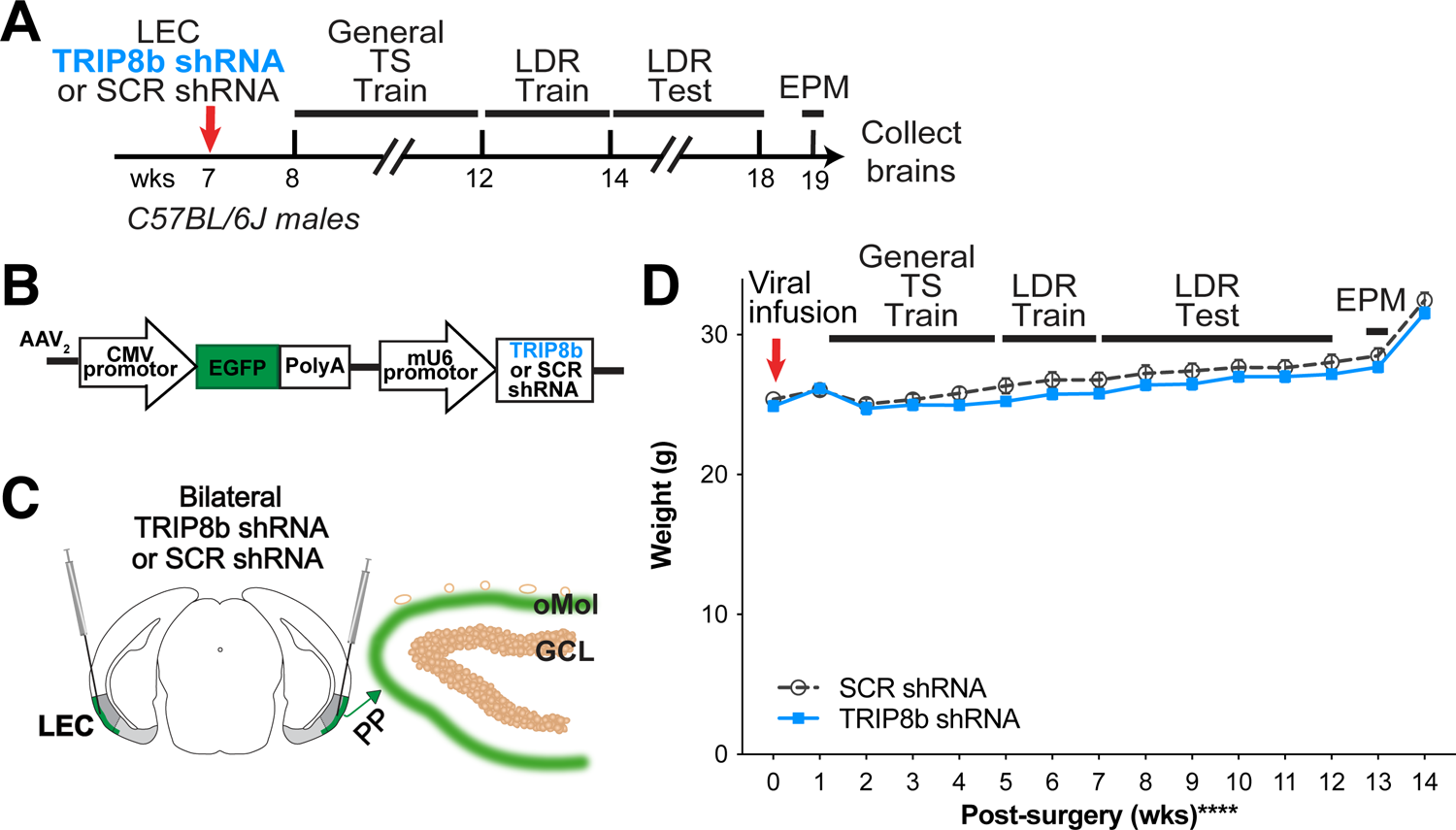
Timeline and overview of experiment. **(A)** Timeline for experiment. Seven-week-old C57BL/6J male mice received bilateral infusions (red arrow) of AAV-TRIP8b short-hairpin (TRIP8b shRNA) or scrambled shRNA (SCR shRNA) into the lateral entorhinal cortex (LEC) to knockdown (KD) the HCN auxiliary protein TRIP8b as done in prior work. After sufficient time for the virus to express encoded proteins, mice went through touchscreen training (General TS Train) with a twelve-window grid (2×6) followed by location discrimination reversal training (LDR Train) and finally testing (LDR Test). After touchscreen experiments were finished, mice were run on the elevated plus maze (EPM) to gauge innate fear of open vs. closed EPM arms. **(B)** Schematic of TRIP8b shRNA construct packaged into adeno-associated virus (AAV_2_) which also expresses EGFP. For AAV-expressing SCR shRNA, the construct was the same with the exception that SCR shRNA (not the TRIP8b shRNA) was under the control of the mU6 promoter. **(C**, left) Schematic of coronal section through an adult mouse brain depicting bilateral LEC AAV infusions (indicated by syringes) given via stereotaxic surgery. Mice received an LEC-directed virus containing either TRIP8b shRNA or a control scrambled (SCR) virus. **(C**, right) Schematic of a portion of an enlarged coronal section through the hippocampal dentate gyrus (DG) ∼4 weeks post-surgery. When processed for immunohistochemistry, “on target” LEC viral infusions would result in expression of fluorescence protein (EGFP+, green) in the LEC projections that course through the perforant path (PP) and terminate in the DG outer molecular layer (oMol, green). **(D)** Body weight in TRIP8b shRNA vs. SCR shRNA mice throughout the experiment shown in **(A)**. Mean+/-SEM. Two-way RM ANOVA was performed in **(D)**. Main effects: Time F (14, 238) = 130.6, ****p<0.0001 and Treatment F (1, 17) = 0.6704, p=0.4242; Interaction: Treatment X Time F (14, 238) = 0.9406, p=0.5158. CMV, cytomegalovirus; GCL, granule cell layer; wks, weeks. Complete statistical information provided in **Supplementary Table 1**.

#### General Touchscreen Training

(prior to LDR) or “General TS Train” **(Fig. 1A)**, consists of five stages (Whoolery et al., 2020; Soler et al., 2021): Habituation, Initial Touch, Must Touch, Must Initiate, and Punish Incorrect (PI). Methods for each stage are described in turn below. Mice went through General TS Train with twelve windows (2×6) for the LDR experiment.

#### Habituation

Mice are individually placed in a touchscreen chamber for 30min (maximum) with the magazine light turned on (LED Light, 75.2 lux). For the initial reward in each habituation session, a tone is played (70 decibel [dB] at 500Hz, 1000ms) at the same time as a priming reward (150ul Ensure® Original Strawberry Nutrition Shake) is dispensed to the reward magazine. After a mouse inserts and removes its head from the magazine, the magazine light turns off and a 10 second (s) delay begins. At the end of the delay, the magazine light is turned on and the tone is played again as a standard amount of the reward (7ul Ensure) is dispensed. If the mouse’s head remains in the magazine at the end of the 10s delay, an additional 1s delay is added. A mouse completes Habituation training after they collect 25 rewards (25 × 7ul) within 30min. Mice that achieve habituation criteria in <30min are removed from the chamber immediately after their 25th reward in order to minimize extinction learning. The measure reported for Habituation is days to completion.

#### Initial Touch

A 2×6 window grid is placed in front of the touchscreen for the remaining stages of training. At the start of the session, an image (a lit white square) appears in a pseudo-random location in one of the 12 windows on the touchscreen. The mouse has 30s to touch the lit square (typically with their nose). If the mouse does not touch the image, it is removed, a reward (7ul Ensure) is delivered into the illuminated magazine on the opposite wall from the touchscreen, and a tone is played. After the reward is collected, the magazine light turns off and a 20s intertrial interval (ITI) begins. If the mouse touches the image while it is displayed, the image is removed and the mouse receives 3 times the normal reward (21ul Ensure, magazine is illuminated, tone is played). For subsequent trials, the image appears in another of the 12 windows on the touchscreen, and never in the same location more than 3 consecutive times. Mice reach criteria and advance past Initial Touch training when they complete 25 trials (irrespective of reward level received) within 30min. Mice that achieve Initial Touch criteria in <30min are removed from the chamber immediately after their 25th trial. The measure reported for Initial Touch is days to completion.

#### Must Touch

Similar to Initial Touch training, an image appears, but now the window remains lit until it is touched. If the mouse touches the lit square, the mouse receives a reward (7ul Ensure, magazine is illuminated, tone is played). If the mouse touches one of the blank windows, there is no response (no reward is dispensed, the magazine is not illuminated, and no tone is played). Mice reach criteria and advance past Must Touch training after they complete 25 trials within 30min. Mice that achieve Must Touch criteria in <30min are removed from the chamber immediately after their 25th trial. The measure reported for Must Touch is days to completion.

#### Must Initiate

Must Initiate training is similar to Must Touch training, but a mouse is now required to initiate the training by placing its head into the already-illuminated magazine. A random placement of the image (lit white square) will then appear on the screen, and the mouse must touch the image to receive a reward (7ul Ensure, magazine lit, tone played). Following the collection of the reward, the mouse must remove its head from the magazine and then reinsert its head to initiate the next trial. Mice advance from Must Initiate training after they complete 25 trials within 30min. Mice that achieve Must Initiate criteria in <30min are removed from the chamber immediately after their 25th trial. The measure reported for Must Initiate is days to completion.

#### Punish Incorrect (PI)

Punish Incorrect training builds on Must Initiate training, but here if a mouse touches a portion of the screen that is blank (does not have a lit white square), the overhead house light turns on and the lit white square disappears from the screen. After a 5s timeout period, the house light turns off and the mouse has to initiate a correction trial where the lit white square appears in the same location on the screen. The correction trials are repeated until the mouse successfully presses the lit white square; however, correction trials are not counted towards the final percent correct criteria. Mice reach criteria and advance past Punish Incorrect training and onto Location Discrimination Reversal Train/Test after they complete 30 trials within 30min at 76% (19 correct) on Day 1 and >80% (>24 correct) on Day 2 over two consecutive days. Mice that achieve PI criteria in <30 min are removed from the chamber immediately after their 30th trial. As with the other stages, a measure reported for PI is days to completion (to reach criteria). However, since the PI stage also contains a metric of accuracy, more measures were analyzed relative to the other five stages. Therefore, other measures reported for PI are session length, trial number, and percent correct responses.

#### Location Discrimination Reversal

(LDR; program LD1 choice reversal v3; ABET II software, Cat #89546-6) tests the ability to discriminate 2 conditioned stimuli that are separated either by a large or small separation. The reversal component of LDR is used here and, as in classic LDR studies (Clelland et al., 2009; Oomen et al., 2013; Swan et al., 2014; Soler et al., 2021), is used to test cognitive flexibility. Taken together, LDR is a hippocampal-dependent task which allows assessment of both discrimination ability as well as cognitive flexibility. In our timeline (**Fig. 1A**), mice received one additional training step (“LDR Train”) prior to the actual 2-choice LDR Test.

#### Location Discrimination Reversal Train (LDR Train)

Mice initiate the trial, which leads to the display of two identical white squares (25 × 25 pixels, **Fig. 3A**) presented with two blank (unlit) squares between them, a separation which is termed “intermediate” (8th and 11th windows in 2×6 high grid-bottom row). One of the left or right locations of the squares is rewarded (i.e. S+) and the other is not (S-), and the initial rewarded location (left or right) is counterbalanced within-group. On subsequent days, the rewarded square location is switched based on the previous day’s performance. Reward side is carried over from the previous session/day, if they did not reach criteria (+1 reversal). A daily LDR Train session is complete once the mouse runs 50 trials or when 30min has passed. Once 7 out of 8 trials had been correctly responded to, on a rolling basis, the rewarded square location was switched (becomes S-), then S+, then S-, etc.; this is termed a “reversal”. Once the mouse reaches >1 reversal in 3 out of 4 consecutive testing sessions, the mouse advances to LDR Test. Measures reported for LDR Train are: percent of each group reaching criteria over time, days to completion, trial number, percent correct to 1st reversal, and time to reach the 1st reversal.

#### Location Discrimination Reversal Test (LDR Test)

Mice initiate the trial, which leads to the display of two identical white squares, either with four black squares between them (“large” separation, two at maximum separation [7th and 12th windows in the bottom row of a 2×6 grid] or directly next to each other (“small” separation, two at minimum separation [9th and 10th windows in the bottom row of a 2×6 grid; **Fig. 4A, B**]). As in LDR Train, only one of the square locations (right-most or left-most) is rewarded (S+, same side for both Large and Small separation, and counterbalanced within-groups). The rewarded square location is reversed based on the previous day’s performance (S+ becomes S-, then S+, then S-, etc). Once 7 out of 8 trials correctly responded to, on a rolling basis, the rewarded square location is reversed (becomes S-, then S+, then S-, etc.). Each mouse is exposed to only one separation type during a daily LDR Test session (either large or small) and the separation type changes every two days (two days of Large, then two days of Small, two days of Large, etc.). A daily LDR Test session is complete once the mouse touches either S+ or S-81 times or when 30min has passed. LDR Test data are analyzed by block (1 block=4 days LDR Test counterbalanced with 2 Large and 2 Small separation daily sessions). Once 24 testing sessions (12 days of Large, 12 days of Small separation) are complete, mice receive a two-week normal feeding prior to extinction testing. Measures reported for LDR Test are all presented for both Large and Small separation: session length, trial number, percent correct during trials to the 1st reversal, number of reversals, number of blank touches (touching an unlit square), reward collection latency, latency to touch the correct image on the last day of the 1st, 4th, and 6th two-day block were reported, and latency to touch the incorrect image (touching the incorrect lit square; does not include blank window touches) on the last day of the 1st, 4th, and 6th two-day block (to allow assessment in the last day in Large or Small separation testing blocks).

#### Elevated Plus Maze (EPM)

The elevated plus maze consists of two open arms (L 67 x W 6 cm) and two closed arms with walls (L 67 x W 6 x H 17 cm, opaque gray Plexiglas walls and black Plexiglas floor, Harvard Apparatus, #760075). At the start of the test, mice are placed on the far end of the open arms and allowed free movement throughout the maze for 5min. The parameters of the EPM (total distance moved, number of entries and duration spent in the open and closed arms) were scored via EthoVisionXT software (Noldus Information Technology) using nose-center-tail tracking to determine position.

### Duration of Experiment

During the five stages of General TS Train, all SCR shRNA and TRIP8b shRNA mice gained touchscreen-related operant learning experience, which was followed by LDR Train and LDR Test. After completing the LDR Test, all mice underwent EPM to gauge innate fear of open vs. closed EPM arms. Brains were collected after the EPM test. Thus, the duration of the experiment (from the beginning of General TS Train to brain collection) was 3 months.

### Brain Collection and Brain Section Preparation

A single cage of mice was brought into the procedure room at a time, and all mice in the cage were decapitated by IACUC approved scissors within 3min. Brains were immersion fixed with 4% paraformaldehyde in 1XPBS at room temperature for 3d followed by cryoprotection (placement in 30% sucrose in 1XPBS at room temperature for 3 more days and then stored at 4°C until sectioning). Brains were sectioned coronally on a microtome (Leica) by covering the brain with fine dry ice and collecting 30µm sections through the entire anterior-posterior length of the hippocampus and entorhinal cortex (distance range from Bregma: −0.82 to −4.24 µm). Serial sets of sections were stored in 0.1% NaN_3_ in 1XPBS at 4°C until processing for slide-mounted immunohistochemistry (IHC) (Ables et al., 2010; Lagace et al., 2010).

### IHC

For single-labeling of tissue with antibodies against either doublecortin (DCX) or GFP, one series of sections was mounted on glass slides (Superfrost/Plus, Fisher) coded to ensure experimenters remained blind throughout quantification and data analysis. Sections were processed for antigen retrieval (0.01 M citric acid, pH 6.0, 95°C, 15 min) and nonspecific staining was blocked by incubating in blocking solution (3% normal donkey serum [NDS], vol/vol in 0.1% Triton X-100 in 1XPBS) for 30min. After blocking, sections were then incubated in either goat anti-DCX (1:5000; Santa Cruz, Cat. #SC-8066) or chicken anti-GFP (1:3000; Aves Cat. #GFP-1020) in 0.1% Tween-20 in 1XPBS overnight. The following day, sections were rinsed and incubated in either biotinylated-donkey anti-goat IgG antibody (Cat. #705-065-003) or biotinylated-donkey-a-chicken-IgY (Cat. #703-065-155), both 1:200 (Jackson ImmunoResearch Laboratories Inc., West Grove, PA) in 1.5% NDS in 1XPBS for 1h. After rinsing in 1XPBS and 30min in 0.3% hydrogen peroxide in 1XPBS to quench endogenous peroxidases, sections were incubated in avidin-biotin complex (ABC Elite, Vector Laboratories) for 60min. After rinsing in 1XPBS, staining for DCX immunoreactive (+) cells was visualized using DAB (Thermo Scientific, Cat. #1856090) and for GFP+ cells was visualized using Fluorescein-labeled Tyramide (PerkinElmer, Cat. #SAT701). Nuclear Fast Red (Vector Laboratories, Cat. #H-3403) or DAPI (Roche, Cat. #236276) were used as counterstains for DCX and GFP immunolabeling, respectively.

### Targeting LEC➔DG Neurons

Accurate virus targeting was verified after brain collection. Specifically, TRIP8b shRNA mice with GFP+ soma in LEC II/III, GFP+ processes in perforant path, GFP+ terminals in the middle and/or outer DG Mol — but with no GFP+ projections in other hippocampal regions, such as CA1 — were considered “on target”. TRIP8b shRNA mice that also had fine projections in non-DG hippocampal regions, such as in the stratum lacunosum-moleculare (SLM), were considered to have too broad of viral expression to be considered on target. Images and functional importance of this targeting is provided in **Supplementary Fig. 1**. No SCR shRNA mice were excluded based on GFP+ processes and terminals since this construct is considered to be biologically inactive (see Discussion).

### Quantification of DCX+ Cells

Unbiased quantification of DCX+ cells (total as well as late progenitors and immature neurons in the superior [or suprapyramidal] and inferior [or infrapyramidal] blade of the DG) was performed via stereology (Latchney et al., 2014; Whoolery et al., 2017; Luna et al., 2019; Clark et al., 2021). The microscope used was a BX51 (Olympus America, Center Valley, PA, USA) with a 40X, 0.63 NA oil-immersion objective. DCX+ cells in the DG subgranular zone (SGZ) were counted. The total DCX+ cell number was calculated by this formula (Yun et al., 2018b; Clark et al., 2021):

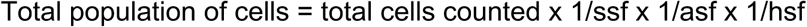

where ssf is the section sampling fraction (DCX: 1/8), asf is the area sampling fraction (1 for these rare populations of cells; thus, all cells were counted in “ssf [e.g. 1/8]” sections), hsf is the height sampling fraction (1 given the minimal effect edge artifacts have in counting soma <10um with ssf 1/8) as described in prior work (Lagace et al., 2010). One hemisphere in hippocampal dorsal DG (−0.95mm ∼ −2. 65mm) was counted for DCX+ cells, thus the resulting formula was:

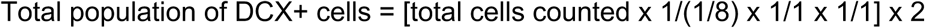

Therefore, the DCX+ cell counts were multiplied by 16 to obtain the total number of DCX+ GC layer (GCL) cells. Similar formulas were applied for DCX+ subtypes (progenitors and immature neurons) which were classified based on their soma morphology and absence/presence of processes.

### Computer Scripts

Prior to statistical analysis, touchscreen data were sorted and extracted. We used a custom Python 3.8.6, and Pycharm script developed by the Eisch Lab to extract, calculate needed values, and compile the data into a database. Extracting the data into an output CSV file was managed with another custom script, and these outputs were verified manually. Following this verification, the data were analyzed using GraphPad Prism 9 according to the tests detailed in the Statistical Analysis section. These scripts along with sample data files are available at github.com/EischLab/18AExtinction.

### Rigor, Additional ARRIVE 2.0 Details, and Statistical Analysis

The experimental unit in this study is a single mouse. For behavioral studies, mice were randomly assigned to groups. Steps were taken at each experimental stage to minimize potential confounds. For example, mice from the two experimental groups (SCR shRNA and TRIP8b shRNA) were interspersed throughout housing racks at CHOP (to prevent effects of cage location) and were interdigitated for all weighing and behaviors (to prevent an order effect). Sample sizes were pre-determined via power analysis and confirmed on the basis of extensive laboratory experience and consultation with CHOP and PennMed statisticians as previously reported (Whoolery et al., 2020; Soler et al., 2021). Exact subject number for each group is provided in **Supplementary Table 1**. A total of n=2 SCR shRNA mice were outliers based on *a priori* established experimental reasons (n=1 SCR shRNA mice did not complete the PI stage even by Day 33; n=1 Sham did not complete LDR “Acquisition” since it reached a humane endpoint) and the data from these mice were excluded from this experiment. The initial subject numbers per group were SCR shRNA (n)=11 and TRIP8b shRNA (n)=16. After assessment for GFP+ terminals restricted to the DG Mol, the subject numbers per group were SCR shRNA (n)=9 and TRIP8b shRNA (n)=10, and these were the mice whose data are presented in the main figures. Experimenters were blinded to treatment until analysis was complete. Data for each group are reported as mean ± s.e.m (line graphs) or median and quartile (violin plot). Testing of data assumptions (normal distribution, similar variation between control and experimental groups, etc.) and statistical analyses were performed in GraphPad Prism. Analyses were hypothesis-based and therefore pre-planned, unless otherwise noted in Results. D-Agostino’s tests and QQ plot were chosen for normality tests. Since the data in all figures passed the normality test, parametric tests were selected. Statistical approaches and results including statistical analysis significance (p-values) and effect size (when RM two-way ANOVA or One-way ANOVA, p<0.05: partial omega-squared wp2 where 0.05 small, 0.06 medium, 0.14 large) are provided in Results and/or **Supplementary Table 1**. Analyses with two groups were performed using an unpaired, two-tailed Student’s t-test. Analyses with more than two variables were performed using two-way ANOVA or Mixed-effects analysis with Bonferroni post hoc test; repeated measures (RM) were used where appropriate, as indicated in figure legends and **Supplementary Table 1**. Analysis of the distribution of subjects reaching criteria between control and experimental groups (survival curve) was performed with the Mantel-Cox test and significance was defined as *p < 0.05.

## RESULTS

### In C57BL/6J male mice, LEC TRIP8b shRNA does not change body weight or influence operant learning on a touchscreen platform

C57BL/6J male mice received bilateral LEC infusions of either SCR shRNA or TRIP8b shRNA one week prior to the start of touchscreen experiments **(Fig. 1A-B)**. This results in GFP+ LEC stellate cells in layers II/III (LECII/III) and GFP+ processes in the perforant path and DG Mol **(Fig. 1C)** (Yun et al., 2018b); mice with GFP+ processes in non-DG regions of the hippocampus were excluded from most data shown in the main text (see below and **Supplementary Fig. 1)**. While our construct can be expressed in non-stellate LEC cells, TRIP8b in the LEC is primarily expressed in stellate cells (Wilkars et al., 2012). Thus, we consider this targeting to result in TRIP8b KD in LECII/III stellate cells and their afferents to the DG Mol, which enhances DG activity-dependent processes and leads to behavior that could be considered “antidepressive-like” (Yun et al., 2018b). SCR shRNA and TRIP8b shRNA mice had similar weight gain throughout the experiment (**Fig. 1D, Supplementary Table 1**), consistent with prior work (Yun et al., 2018b). In addition, SCR shRNA and TRIP8b shRNA mice completed each stage of the initial operant touchscreen training in similar periods of time **(Fig. 2A, Supplementary Table 1)**. During PI, both SCR shRNA and TRIP8b shRNA mice showed high variability in days to completion. Therefore, we examined additional PI parameters on Day 1 vs. Last Day. SCR shRNA and TRIP8b shRNA mice had similar PI session length **(Fig. 2B, Supplementary Table 1)** and number of trials **(Fig. 2C, Supplementary Table 1)**. However, there was a main effect of Day in two-way RM ANOVA for the variable of percent correct **(Supplementary Table 1)**. Both SCR shRNA and TRIPb8b shRNA mice had higher accuracy on PI Last Day vs. Day 1 **(Fig. 2D, Supplementary Table 1)**. Thus, SCR shRNA and TRIP8b shRNA mice perform similarly in the fundamental operant learning that is required on this operant touchscreen platform, improving accuracy across the duration of the PI training stage.

**Figure 2.**
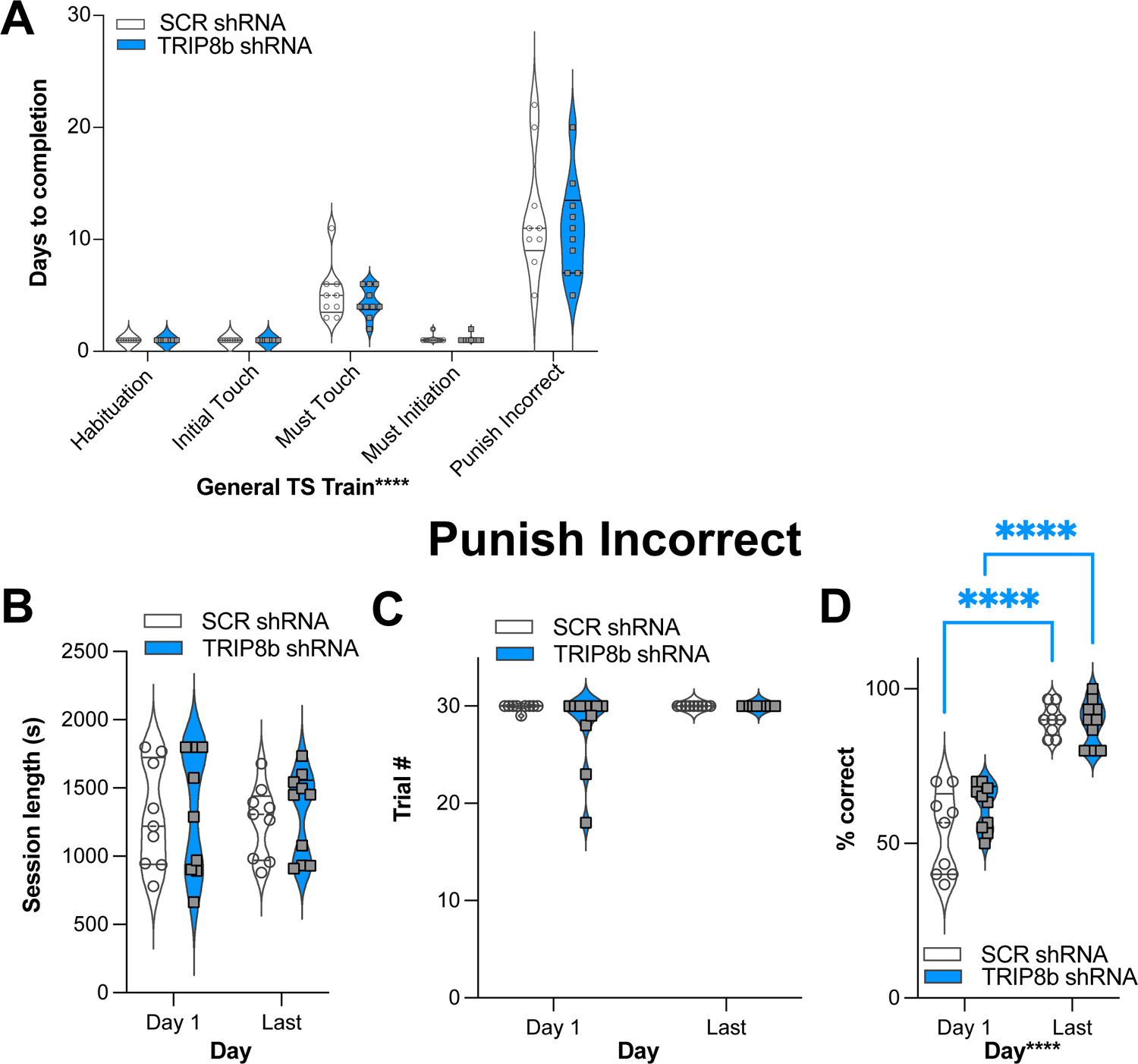
Knockdown of TRIP8b in the LEC does not change performance in general TS Train. **(A)** In the five stages of General TS Train (Habituation, Initial Touch, Must Touch, Must Initiate, and Punish Incorrect), SCR shRNA and TRIP8b shRNA mice took a similar number of days to reach criteria. **(B-D)** In other criteria (beyond Days) for the Punish Incorrect stage of General TS Train, SCR shRNA and TRIP8b shRNA mice also performed similarly on both the first and last day: session length **(B)**, number of trials **(C)**, and % correct (accuracy). **(D)**. Note both SCR and TRIP8b mice showed increased accuracy between the first and last day of Punish Incorrect. Two-way RM ANOVA was performed in **(A-D)**: **(A)** Main Effects: Training Stage F (4, 68) = 75.03, ****p<0.0001 and Treatment F (1, 17) = 0.6292, p=0.4386; Interaction: Training Stage x Treatment F (4, 68) = 0.3333, p=0.8547. **(B)** Main effects: Training Stage F (1, 17) = 0.08579, p=0.7731 and Treatment F (1, 17) = 0.2492, p=0.6240; Interaction: Training Stage X Treatment F (1, 17) = 2.128e-005, p=0.9964. **(C)** Main effects: Training Stage F (1, 17) = 2.858, p=0.1092 and Treatment F (1, 17) = 2.335, p=0.1449; Interaction: Training Stage X Treatment F (1, 17) = 2.335, p=0.1449. **(D)** Main Effects: Training Stage F (1, 17) = 148.2, ****p<0.0001 and Treatment F (1, 17) = 1.599, p=0.2231, *post hoc*: ****p<0.0001 in Day 1 vs Last Day in both SCR and TRIP8b; Interaction: Training Stage X Treatment F (1, 17) = 3.358, p=0.0844. LEC, lateral entorhinal cortex; s, seconds; S+, stimulus associated with a reward; TS, touchscreen. Complete statistical information provided in **Supplementary Table 1**.

### LEC TRIP8b shRNA mice — but not SCR shRNA mice — have better discrimination accuracy on Day 1 vs. the last day of LDR Train

Having established similar operant learning in SCR mRNA and TRIP8b shRNA mice, we next assessed their performance in LDR Train **(Fig. 3A)**, the precursor to LDR Test where lit squares are separated by an intermediate number of unlit squares. There was a visual difference in percent of SCR shRNA and TRIP8b shRNA subjects reaching the criteria in LDR Train (perhaps due to one mouse — which was not a statistical outlier — in TRIP8b group taking 15 days), but this difference was rejected by survival curve analysis **(Fig. 3B, Supplementary Table 1)**. SCR shRNA and TRIP8b shRNA mice also had similar average days to complete LDR Train **(Fig. 3C, Supplementary Table 1)**. Interestingly, TRIP8b shRNA mice had higher accuracy on the last day of LDR Train vs. Day 1, while SCR shRNA mice had similar accuracy between the last day and Day 1 **(Fig. 3D, Supplementary Table 1)**. However, SCR shRNA and TRIP8b shRNA mice did not differ in total session time to reach the 1st reversal **(Fig. 3E, Supplementary Table 1)** or % correct to 1st reversal **(Fig. 3F, Supplementary Table 1)**. Thus, in the training step for LDR, LEC TRIP8b shRNA mice — but not LEC SCR shRNA mice — have better discrimination accuracy on Day 1 vs. the last day.

**Figure 3.**
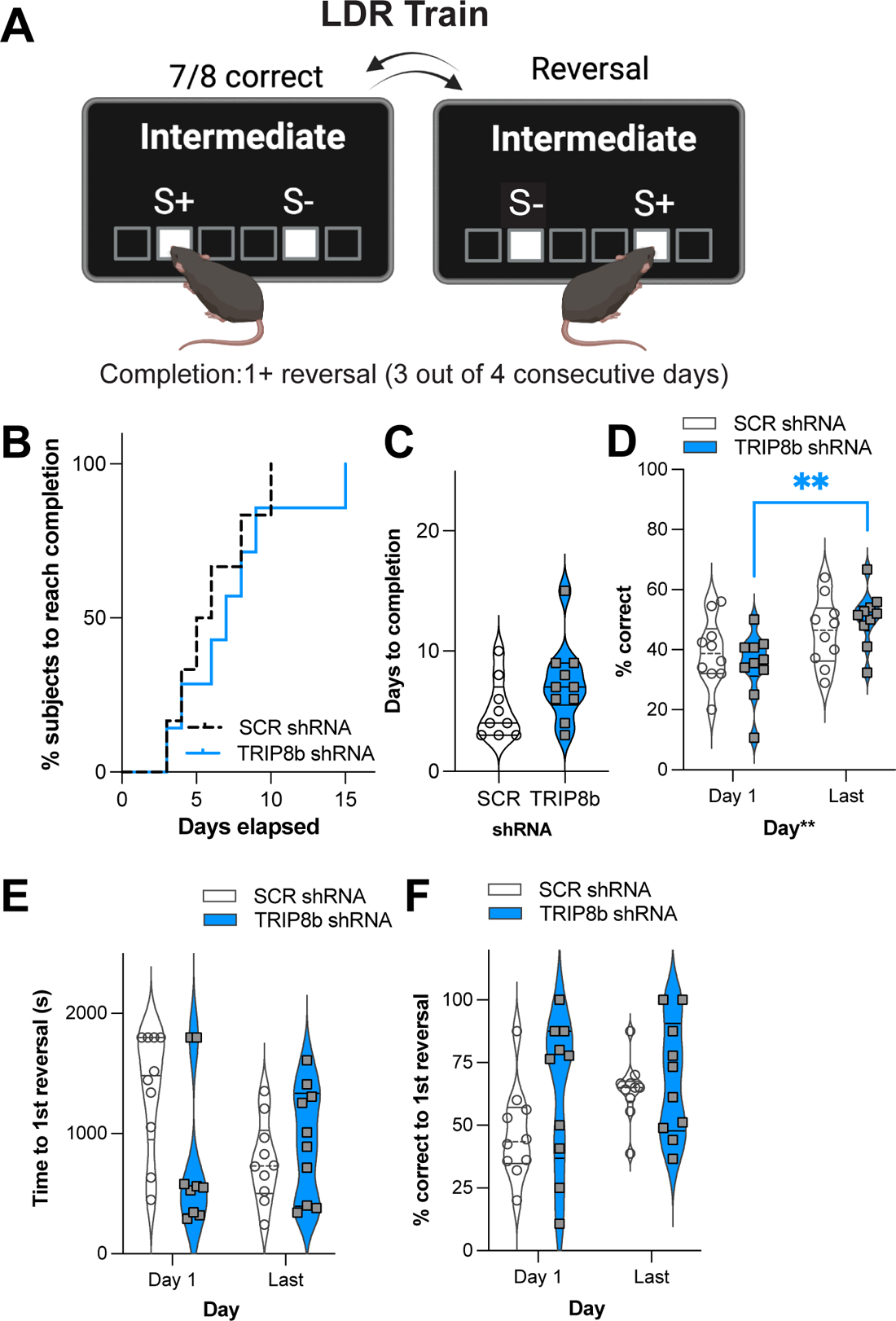
Knockdown of TRIP8b in the LEC improved accuracy between Day 1 vs. Last Day of LDR Train. **(A)** Schematic depicting LDR Train sessions, criteria for reversal, and criteria for LDR Train completion. When mice made 7 out of 8 consecutive correct choices of stimulus (S+), a reversal occurred (S+ became S-[incorrect stimulus] and previous S- became S+). Criteria for LDR Train completion is a mouse making at least 1 (1+) reversal on 3 out of 4 consecutive days. **(B-C)** LDR Train data from SCR shRNA and TRIP8b shRNA mice showing **(B)** % subjects that complete LDR Train over days elapsed and **(C)** total days it took each group to complete LDR Train. **(D)** Accuracy (% correct) was the same for SCR shRNA mice on LDR Train Day 1 vs. Last Day. In contrast, accuracy was improved for TRIP8b shRNA mice on LDR Train Day 1 vs. Last Day. **(E-F)** On LDR Train Day 1, SCR shRNA and TRIP8b shRNA mice took the same amount of time to reach the 1st reversal **(E)** and had similar accuracy **(F)**. On the last day, there was no difference in speed to first reversal or accuracy between the groups. Mantel-Cox test in **(B)**, p=0.5328, unpaired t-test in **(C)**, p=0.1088 and Two-way RM ANOVA in **(D-F)** were performed. **(D)** Main effects: Time F(1, 17)=10.08, **p=0.0055 and Treatment F(1, 17)=0.0006715, p=0.9796, *post hoc*: **p=0.0075 on Day 1 vs Last Day in TRIP8b; Interaction: Treatment x Time F(1, 17)=2.084, p=0.1671. **(E)** Main effects: Time F(1, 17)=1.386, p=0.2552 and Treatment F(1, 17)=1.500, p=0.2373; interaction: Treatment x Time F(1, 17)=2.382, p=0.1411. **(F)** Main effects: Time F(1, 17)=1.571, p=0.2271 and Treatment F(1, 17)=2.700, p=0.1187; Interaction: Treatment x Time F(1, 17)=0.5309, p=0.4762. LDR, location discrimination reversal; s, seconds. Complete statistical information provided in **Supplementary Table 1**.

### LEC TRIP8b shRNA mice have improved behavioral pattern separation and cognitive flexibility compared to SCR shRNA mice

After mice reached the LDR Train criteria, mice underwent the LDR Test to assess behavioral pattern separation and cognitive flexibility (Mar et al., 2013; Oomen et al., 2013; Phillips et al., 2018). During the LDR Test, various loads with two different spacing between S+ and S- were applied: Large separation (lit squares separated by four unlit squares) and Small separation (lit squares adjacent to each other; **Fig. 4A-H**). Mice in one group were randomly assigned to either Large or Small separation for the first 2 days, and mice in the other group were counterbalanced. As LDR Test consists of 6 blocks (a “block” is 2 days of Large and 2 days of Small separation) over 24 days, behavioral pattern separation and reversal learning can be assessed in early and late LDR Test.

**Figure 4.**
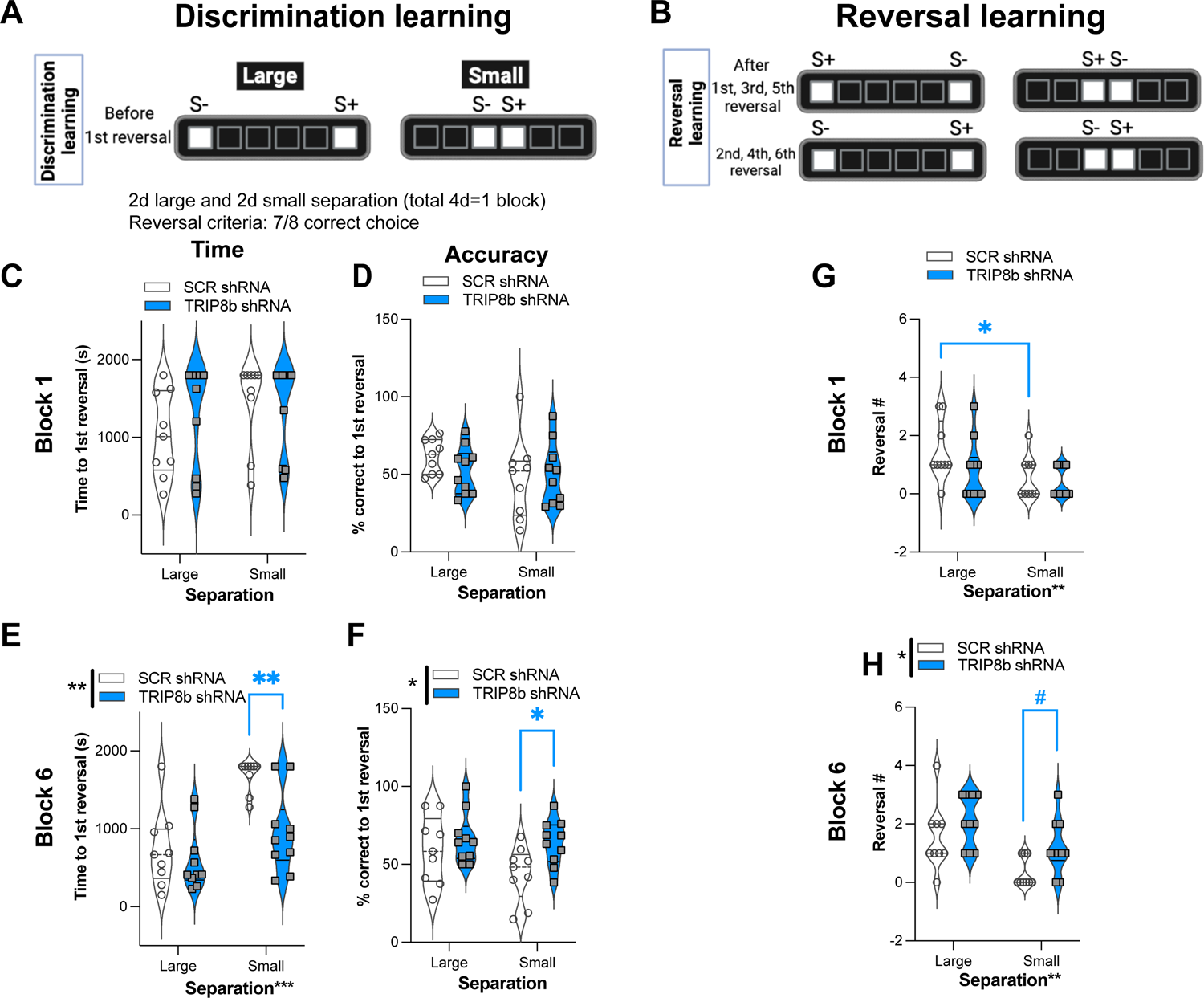
Knockdown of TRIP8b in the LEC improved discrimination learning in LDR Test. **(A-B)** Schematic depicting LDR test sessions. **(A)** Discrimination learning was tested in the sessions that occurred prior to the first reversal each day, with the degree of difficulty provided by the lit square (stimuli, S) being separated by either four black squares (Large separation) or zero spaces (Small separation). After selecting the correct choice (S+) in 7 out of 8 consecutive trials, S+ became S- and the Reversal learning trials **(B)** began, and continued with S+ and S- designated as depicted. LDR Test was run for 6 blocks, where 1 block consisted of 2 days of Large separation and 2 days of Small separation. **(C-F)** Discrimination Learning. **(C, D)** TRIP8b shRNA and SCR shRNA mice performed with similar speed and accuracy before 1st reversal under both Large and Small separation on the last day of the Block 1. **(E, F)** On the last block (Block 6), TRIP8b mice reached the 1st reversal faster and with higher accuracy than SCR shRNA mice, particularly under the condition of Small separation (but not Large) separation. **(G-H)** While TRIP8b shRNA and SCR shRNA mice made a similar number of reversals in Block 1, there is a main effect of Treatment and Separation in Block 6. SCR shRNA mice made fewer reversals under Small vs. Large separation, while TRIP8b shRNA mice made similar reversal numbers under Small and Large separations. Two-way RM ANOVA was performed in **(C-F). (C)** Main effects: Separation F(1,17)=1.084, p=0.3124 and Treatment F(1,17)=2.423, p=0.1380; interaction: Separation x Treatment F(1,17)=0.1799, p=0.6768. **(D)** Main effects: Separation F(1,17)=1.679, p=0.2123 and Treatment F(1,17)=0.3233, p=0.5771; interaction: Separation x Treatment F(1,17)=0.8583, p=0.3672. **(E)** Main effects: Separation F(1,17)=16.36, ***p=0.0008 and Treatment F(1,17)=15.53, **p=0.0011, *post hoc*: **p=0.0012 in SCR vs. TRIP8b on Small separation; interaction: Separation x Treatment F(1,17)=3.589, p=0.0753. **(F)** Main effects: Separation F(1,17)=1.760, p=0.2022 and Treatment F(1,17)=8.939, **p=0.0082, *post hoc*: *p=0.0397 in SCR vs TRIP8b on Small Separation; interaction: Separation x Treatment F(1,17)=0.8945, p=0.3575. **(G)** Main effects: Separation F(1,17)=10.07, **p=0.0056 and Treatment F(1,17)=1.448, p=0.2454; interaction: Separation x Treatment F(1,17)=1.448, p=0.2453, *post hoc*: *p=0.0156 in Large vs. Small separation of SCR shRNA. **(H)** Main effects: Separation F(1,17)=14.83, **p=0.0013 and Treatment F(1,17)=5.531, *p=0.0310, *post hoc*: #p=0.081 in SCR vs. TRIP8b in Small separation; interaction: Separation x Treatment F(1,17)=0.3419, p=0.5664. LDR, location discrimination reversal; s, seconds, S+=stimulus associated with a reward. Complete and detailed statistical information provided in **Supplementary Table 1**.

To examine behavioral pattern separation ability, the total time to reach to 1st reversal and percent correct to 1st reversal were measured in both groups with Large and Small separation on Block 1 (first block, early LDR Test) and Block 6 (last block, late LDR Test; **Fig 4A-F**). In Block 1, SCR shRNA and TRIP8b shRNA took a similar amount of time to reach the 1st reversal **(Fig. 4C, Supplementary Table 1)** with similar accuracy **(Fig. 4D, Supplementary Table 1)** in both Large and Small separation. In Block 6, SCR shRNA and TRIP8b shRNA mice also took a similar amount of time to the first reversal in Large separation. However, in Block 6 Small separation, TRIP8b shRNA mice took 43% less time vs. SCR shRNA mice to reach the first reversal **(Fig. 4E, Supplementary Table 1)**. Moreover, while SCR shRNA and TRIP8b shRNA mice performed with similar accuracy in Block 6 Large separation, in Block 6 Small separation TRIP8b shRNA mice were 45% more accurate vs. SCR shRNA **(Fig. 4F, Supplementary Table 1)**. These data suggest in late LDR Test, LEC TRIP8b KD enhances behavioral pattern separation when stimuli locations are challenging to differentiate, and thus cognitive load is high (Yassa et al., 2011; Bekinschtein et al., 2013; Kent et al., 2015a, 2015b; Kassab and Alexandre, 2018).

### LEC TRIP8b shRNA mice have improved cognitive flexibility compared to SCR shRNA mice

In LDR Test, early and late cognitive flexibility can be inferred by the number of reversals made in Block 1 **(Fig. 4G)** and Block 6 **(Fig. 4H)**, respectively. In Block 1, SCR shRNA mice made fewer reversals in Small separation vs. Large separation **(Fig. 4G, Supplementary Table 1)**, reflecting the difficulty of Small separation (high load) vs. Large separation (low load). However, in Block 6, TRIP8b shRNA mice accomplished 260% more reversals in Small — but not Large — separation vs. SCR shRNA mice **(Fig. 4H)**. This suggests LEC TRIP8b KD improves cognitive flexibility under high load.

### Improved behavioral pattern separation is seen in mice where TRIP8b shRNA is expressed in LEC➔DG Mol neurons but not in LEC➔DG Mol + CA1 SLM neurons

Axon terminals from stellate cells in the LEC and MEC terminate in the outer Mol of DG and also in non-DG regions, such as the SLM of CA1, respectively (Witter, 2007; Kohara et al., 2014; Kitamura et al., 2015; Witter et al., 2017; Vandrey et al., 2020). All TRIP8b shRNA data presented up to this point reflect only “on target” mice where the expression of GFP+ was largely restricted to LEC➔DG Mol neurons: GFP+ cell bodies in the LEC layer IIa (largely excluded from layer IIb) and GFP+ processes and terminals in DG Mol, but not in CA1 SLM (**Supplementary Fig. 1B, D**). To assess if the TRIP8b shRNA-induced improvement in behavioral pattern separation and cognitive flexibility was restricted to only these “on target” LEC➔DG Mol mice, we compared behavioral output among TRIP8b shRNA LEC➔DG Mol, SCR shRNA mice, and TRIP8b shRNA LEC➔DG Mol+CA1 mice (where GFP+ cell bodies were seen in LEC IIb and GFP+ processes and terminals were seen in SLM CA1; **Supplementary Fig. 1C-D**). In Block 6 under conditions of Large separation, measures of behavioral pattern separation (Time to 1st reversal, % correct to 1st reversal, **Supplementary Fig. 1E-F**) and cognitive flexibility (number of reversals achieved, **Supplementary Fig. 1G**) were similar among all three groups of mice (SCR shRNA, TRIP8b shRNA LEC➔DG Mol, and TRIP8b shRNA LEC➔DG Mol+CA1 mice). However, in Block 6 under conditions of Small separation, TRIP8b shRNA LEC➔DG Mol mice reached the 1st reversal faster (**Supplementary Fig. 1H**) and had greater accuracy to the 1st reversal (**Supplementary Fig. 1I**) compared to SCR shRNA mice, suggesting improved behavioral pattern separation. There was a visual difference in these measures of behavioral pattern separation between SCR shRNA mice and TRIP8b shRNA LEC➔DG Mol+CA1 mice, but the difference was rejected by one-way ANOVA. In contrast to the improved behavioral pattern separation in TRIP8b shRNA LEC➔DG Mol mice vs. SCR shRNA mice, a parameter of cognitive flexibility (number of reversals) was not different among the three groups of mice (**Supplementary Fig. 1J**).

### The improved behavioral pattern separation and cognitive flexibility seen in LEC TRIP8b shRNA mice are not a reflection of altered attention, motivation, or impulsivity

Circuit-based manipulations can influence LDR Test performance by indirectly changing attention, motivation, or impulsivity. To test this possibility, data were extracted from the LDR Test sessions to assess latency to select a correct image, latency to collect the reward from the food hopper, and total number of blank touches made during the ITI which are measures of attention, motivation and impulsivity, respectively. These LDR Test measures were collected from both early (Block 1) and late (Block 6) and under Large and Small separation conditions (**Fig. 5**). Analysis revealed no difference between SCR shRNA and TRIP8b shRNA mice in any metric or condition (**Fig. 5A-F**). Thus, changes in attention, motivation, or impulsivity do not contribute to the improved behavioral pattern separation and cognitive flexibility seen in TRIP8b shRNA vs. SCR shRNA mice.

**Figure 5.**
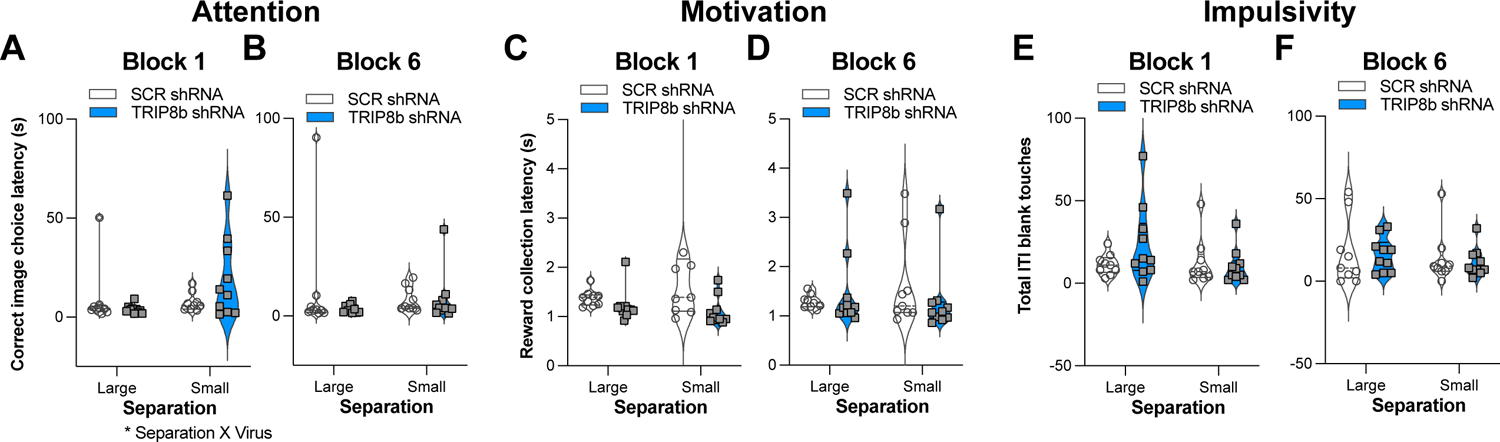
Knockdown of TRIP8b in the LEC did not change measures relevant to attention, motivation or impulsivity during the LDR Test. **(A-F)** TRIP8b shRNA mice and SCR shRNA mice performed similarly on the following measures collected during Block 1 **(A, C, E)** or Block 6 **(B, D, F)** when faced with a stimuli separation that was either Large or Small in LDR Test: **(A, B)** Latency to choose the correct stimulus; **(C, D)** Latency to collect the reward from the food hopper; and **(E, F)** Number of touches to a blank screen during the inter-trial interval (ITI). Two-way RM ANOVA for all panels. **(A)** Main effects: Separation F(1,17)=2.977, p=0.1026 and Treatment F(1,17)=0.4937, p=0.4918, *post hoc*: p=0.1204 in SCR vs TRIP8b on Small separation; Interaction: Separation x Treatment F(1,17)=4.610, *p=0.0465. **(B)** Main effects: Separation F(1,17)=0.0003390, p=0.9855 and Treatment F(1,17)=0.8853, p=0.3599; Interaction: Separation x Treatment F(1,17)=0.8399, p=0.3722. **(C)** Main effects: Separation F(1,17)=1.144, p=0.2997 and Treatment F(1,17)=2.248, p=0.1521; Interaction: Separation x Treatment F(1,17)=1.539, p=0.2316. **(D)** Main effects: Separation F(1,17)=0.09015, p=0.7676 and Treatment F(1,17)=0.1240, p=0.7291; Interaction: Separation x Treatment F(1,17)=1.383, p=0.2559. **(E)** Main effects: Separation F(1,17)=3.077, p=0.0974 and Treatment F(1,17)=0.7417, p=0.4011, *post hoc: p=0*.*1537* in SCR vs TRIP8b on large separation; Interaction: Separation x Treatment F(1,17)=4.547, *p=0.0478. **(F)** Main effects: Separation F(1,17)=1.877, p=0.1885 and Treatment F(1,17)=0.08871, p=0.7694; Interaction: Separation x Treatment F(1,17)=0.0446, p=0.8353. LDR, location discrimination reversal; s, seconds. Complete statistical information provided in **Supplementary Table 1**.

### LEC TRIP8b shRNA and SCR shRNA mice have similar exploration and performance in a task of innate anxiety

Another indirect way that circuit-based manipulations can influence LDR Test performance is by changing measures relevant to exploration and innate anxiety. For example, perhaps TRIP8b shRNA mice have a shorter time to 1st reversal in LDR Test because they are also less anxious. This possibility was tested by assessing mice in the EPM (**Fig. 1A**). Consistent with prior work (Yun et al., 2018b), TRIP8b shRNA and SCR shRNA mice moved a similar total distance (**Fig. 6A**), spent a similar amount of time in the open EPM arms (**Fig. 6B**), and entered the open arms at a similar frequency (**Fig. 6C**). These data suggest TRIP8b shRNA-induced changes in innate exploration and anxiety do not contribute to TRIP8b-induced improvement in behavioral pattern separation and cognitive flexibility.

**Figure 6.**
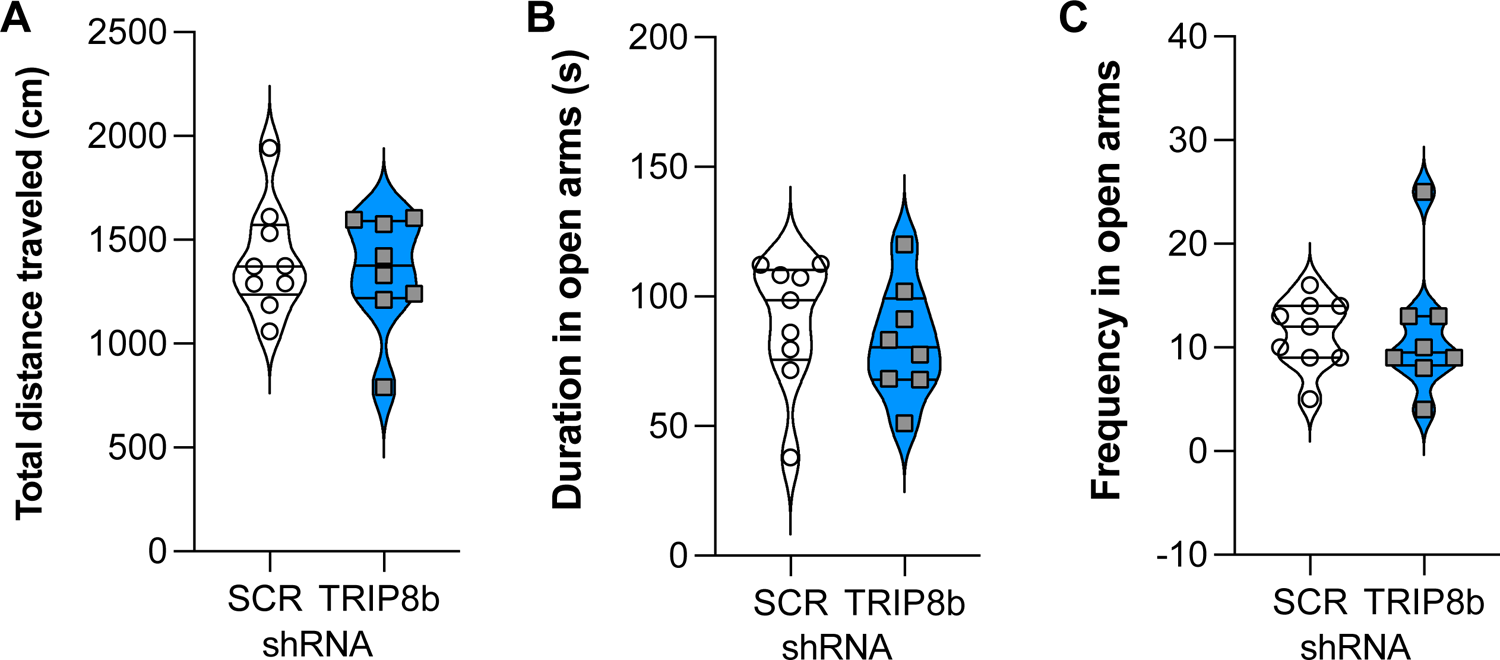
Knockdown of TRIP8b in the LEC does not induce anxiolytic-like behavior. **(A)** SCR shRNA mice and TRIP8b shRNA mice explore similarly in a novel environment, as based on a total movement traveled. **(B-C)** SCR shRNA mice and TRIP8b shRNA mice have similar performance in the elevated plus maze, as based on time spent in open arms **(B)** and frequency to enter to open arms **(C)**. Unpaired t-test for all panels: **(A)** p=0.6529, **(B)** p=0.5044, **(C)** p=0.9863. Complete statistical information provided in **Supplementary Table 1**.

### LEC TRIP8b shRNA mice have more DG neurogenesis compared to SCR shRNA mice

Manipulations that increase EC activity, including EC TRIP8b KD, increase indices of DG neurogenesis (Stone et al., 2011; Yun et al., 2018b). As increased DG neurogenesis is linked to improved behavioral pattern separation and cognitive flexibility (Bekinschtein et al., 2011; Burghardt et al., 2012; Kent et al., 2015c; McAvoy and Sahay, 2017), we hypothesized that LEC TRIP8b shRNA mice would have more DG neurogenesis compared to SCR shRNA mice. The brains from a random subset of mice that underwent behavioral assessment (**Fig. 1A**) were assessed for the number of DCX+ cells in the dorsal DG, and those DCX+ cells were categorized via morphology as progenitor cells or immature neurons (**Fig. 7A-B**). Compared to SCR shRNA mice, TRIP8b shRNA mice had ∼22% more total DCX+ cells and 20% more immature neurons, with no change in the number of progenitors (**Fig. 7C**). We also considered the DCX+ cell counts in the superior vs. inferior blades of the DG as there are functional differences between these aspects of the DG GCL (Collins et al., 2009; Jinno, 2011; Raber et al., 2011; Alves et al., 2018; Raven et al., 2019). Compared to SCR shRNA mice, TRIP8b shRNA mice had 16% and 30% more DCX+ cells in the superior and inferior blades, respectively (**Fig. 7D**). Based on morphology and compared to SCR shRNA mice, TRIP8b shRNA mice had ∼38% more DCX+ progenitor cells only in the inferior blade (**Fig. 7E**), and 17% and ∼22% more DCX+ immature neurons in the superior and inferior blades, respectively (**Fig. 7F**).

**Figure 7.**
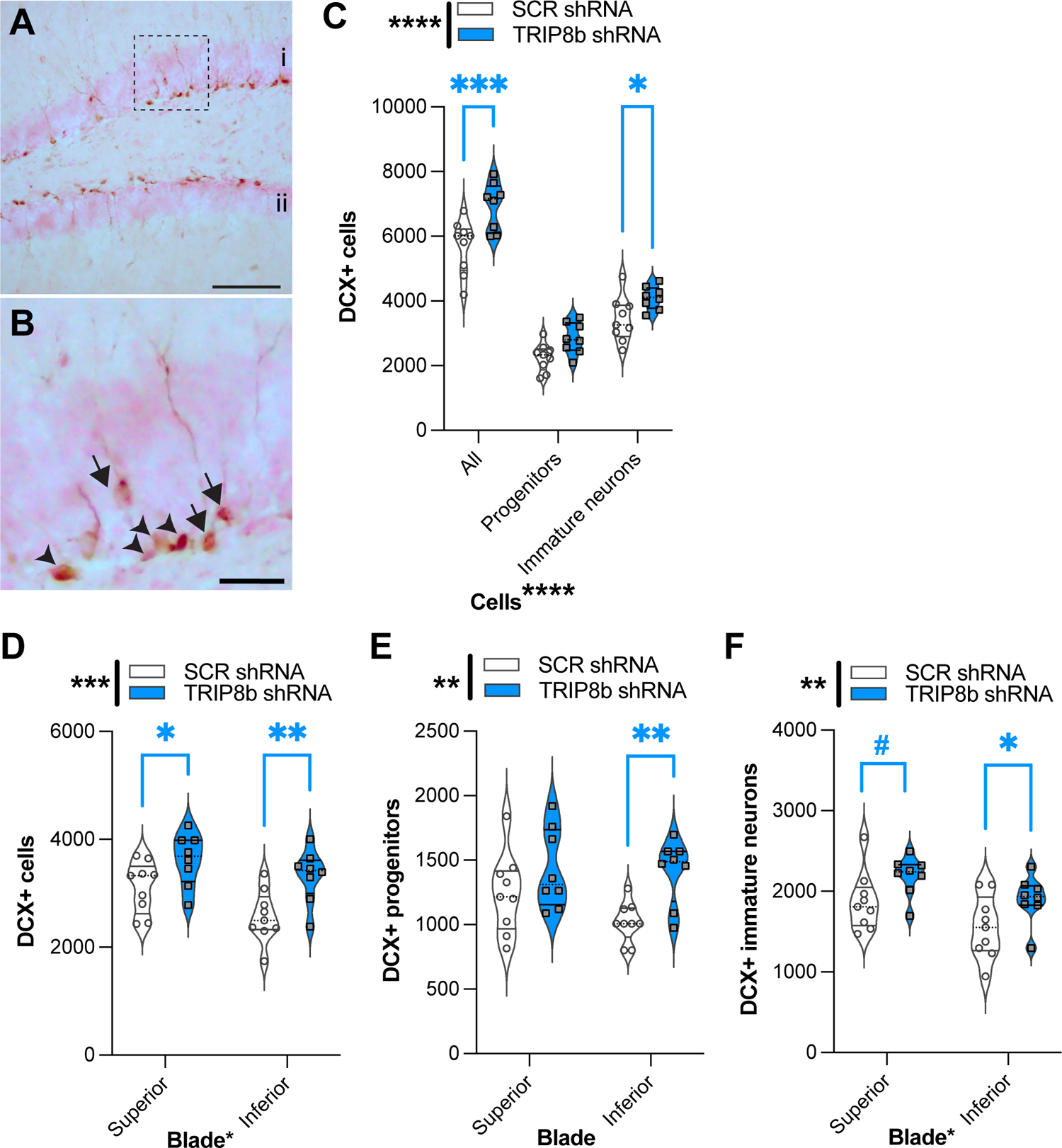
Knockdown of TRIP8b in the LEC increases DCX-immunoreactive (+) cells, an index of dentate gyrus adult neurogenesis. **(A-B)** Representative microscopic images of DCX cells from dorsal DG (i: superior blade, ii: inferior blade in **A**). **(B)** High magnification images for cell types from dotted square in A (Arrow heads: progenitor cells and arrows: immature cells) **(C)** TRIP8b KD in the LEC increases the number of total DCX+ cells and DCX+ immature neurons in the dorsal DG (−0.95 to −2.65mm). **(B)** TRIP8b KD in the LEC increases DCX+ immature neurons in both the suprablade and infrablade of the DG. **(C)** TRIP8b KD in the LEC significantly increases the number of DCX+ late progenitors in the infrablade. Two-way ANOVA for all panels: Main Effects: **(C)** Virus F (1, 45) = 23.58, ****p<0.0001, post hoc: *p=0.0292 in SCR vs. TRIP8b in immature neurons, and Cell type F (2, 45) = 165.4, ****p<0.0001; interaction: Virus X Cell type F (2, 45) = 1.466, p=0.2416. **(D)** Virus F (1, 30) = 14.50, ****p=0.0006, post hoc: *p=0.0237 in SCR vs. TRIP8b in superior, **p=0.0032 SCR vs. TRIP8b in inferior; and region F (1, 30) = 6.466, *p=0.0164 interaction: Virus X Region F (1, 30) = 0.5293, p=0.4726. **(E)** Virus F (1, 30) = 10.55, **P=0.0029, post hoc: **p=0.0041 in SCR vs. TRIP8b in the inferior region, and region F (1, 30) = 1.712, P=0.2006; interaction: Virus X Region F (1, 30) = 1.330, p=0.2579. **(F)** Virus F (1, 30) = 8.813, **P=0.0058, post hoc: #p=0.0548 in SCR vs. TRIP8b in the superior region, *p=0.0357 in SCR vs. TRIP8b in the inferior region, and region F (1, 30) = 7.121, P=0.0122; interaction: Virus X Region F (1, 30) =0.0202, p=0.8878. Scale bars: 100um **(A)**, 25 um **(B)**. Complete statistical information provided in **Supplementary Table 1**.

## DISCUSSION

Increased understanding of the cell- and circuit-based function of LEC➔DG projections will provide insight into many aspects of biomedical research, including cognitive decline in normal aging, disease progression in Alzheimer’s disease, and neurostimulation to combat cognitive dysfunction or other neuropsychiatric symptoms (Albert, 1996; Stranahan and Mattson, 2010; Shin et al., 2014; Zhou et al., 2016; Yu et al., 2019; Igarashi, 2022). Prior to this current work, few studies had examined how projections from the EC specifically to the hippocampal DG subregion — for example, LEC➔DG projections — influence behavior. Here we focused on LEC fan cells for three reasons: they are the only LEC glutamatergic neuron type that directly projects to the DG, they synapse on DG neurons critical for discrimination and reversal learning (e.g. granule and adult-generated neurons), and they are responsible for formation of complex associations required for these cognitive abilities. An ablation study showed that LEC fan cells to the DG are required to discriminate novel object-place-context configurations, but not novel object or object-context recognition in male mice (Vandrey et al., 2020). Also, a lesion study showed that an intact LEC is required for optimal performance in the LDR behavioral pattern separation task (Vivar et al., 2012). As prior work on the function of LEC➔DG projections employed ablation and lesion strategies, here we asked for the first time if the <u>activity</u> of LEC➔DG projections influences behavioral pattern separation in male mice. We also probed, also for the first time, if LEC➔DG projection activity regulated cognitive flexibility. We hypothesized that increased activity of LEC➔DG neurons would improve behavioral pattern separation and reversal learning as EC-DG stimulation does with hippocampal-dependent associative learning. We found that relative to control mice, male mice that received LEC TRIP8b shRNA showed improved indices of both behavioral pattern separation and cognitive flexibility; *post-mortem* analysis showed they also had more DG neurogenesis, supporting that our manipulation increased DG activity as shown in prior work (Yun et al., 2018b). Our findings clarify the circuitry engaged in these higher cognitive abilities, and are novel in that they implicate the activity of the “upstream” LEC in behavioral pattern separation and cognitive flexibility.

There are three particularly notable aspects of the data presented here. First, long-term stimulation of LECIIa fan cells improves measures of location discrimination when the load on pattern separation is high (lit squares are close together) but not when the load is low (lit squares are far apart). Testing discrimination under different loads is a defining feature of behavioral pattern separation tasks (Rolls and Kesner, 2006). This finding was expected; the DG is critical for behavioral pattern separation (Leutgeb et al., 2007; Bakker et al., 2008; Clelland et al., 2009) and increased activity of the EC➔DG pathway improves DG-dependent behavioral tasks (Stone et al., 2011; Xia et al., 2017; Yun et al., 2018b). However, this finding is important. In showing for the first time that increased activity of the LEC➔DG circuit improves behavioral pattern separation, this finding suggests DG dysfunction may be reversed by targeting the activity of afferent regions rather than targeting the DG itself (Suthana et al., 2012; Jacobs et al., 2016).

Second, long-term stimulation of LECIIa fan cells also improves measures of reversal learning. Cognitive flexibility is typically considered to be regulated by the activity and integrity of the prefrontal cortex (PFC) (Kehagia et al., 2010; Izquierdo et al., 2017; Girotti et al., 2018). However, many studies point to a role for the dentate gyrus in cognitive flexibility (Burghardt et al., 2012; Swan et al., 2014; Lucassen and Oomen, 2016; Anacker and Hen, 2017; Berdugo-Vega et al., 2021; Gomes-Leal, 2021). For example, ablation of dentate gyrus adult-generated neurons decreases cognitive flexibility in the location discrimination reversal task (Swan et al., 2014). However, our data are the first to show a role for the activity of the entorhinal cortex in cognitive flexibility. While the specific mechanism underlying this effect is not tested in the present work, one consideration for future research is that induced stimulation of LECIIa fan cells (which project to the DG) may indirectly alter the activity of LEC cells that project to the PFC, such LECII pyramidal cells or neurons in deep LEC layers, or the activity of MEC cells (Ohara et al.; Insausti et al., 1997; Delatour and Witter, 2002; Tanji and Hoshi, 2008; Canto and Witter, 2012; Yu et al., 2021).

A final notable aspect of our data is the timing of the effect; long-term stimulation of LECIIa fan cells improves indices of behavioral pattern separation and cognitive flexibility in late LDR test trials (Blocks 5-6), not in early ones and not during LDR training or general touchscreen training trials. It is unclear why this is the case. One possible answer comes from prior work where inducible ablation of postnatal DG neurogenesis (which is essentially a “long-term” manipulation) impaired cognitive flexibility in late, but not early, trials in the LDR task (Swan et al., 2014). Therefore, it is possible that increased activity of the LEC➔DG circuit increases DG neurogenesis which, over time, leads to compensatory improved function of the DG in behavioral pattern separation and cognitive flexibility tasks. Testing this hypothesis is outside the scope of this work. It warrants pointing out, though, that long-term stimulation of the LEC➔DG circuit likely leads to many compensatory circuit changes, much like ablation of cells/circuits as has been done in prior work (Vivar et al., 2012; Swan et al., 2014; Vandrey et al., 2020). While long-term, but not acute, EC stimulation improves DG-dependent behavior (Yun et al., 2018b), it remains to be tested if a single stimulation followed by time improves DG-dependent behavior, as shown in prior work with manipulation of neurogenesis (Airan et al., 2007; Xia et al., 2017). Alternatively, this could be tested with a behavioral pattern separation task that can be performed over a few days (spontaneous location recognition or object lure discrimination and mnemonic discrimination testing) rather than a month (the present work). Using a more condensed behavioral pattern separation task would also enable future dissociation of the stages of learning (encoding, consolidation, and retrieval) and factors that regulate each stage (Bekinschtein et al., 2013; Johnson et al., 2018; Morales et al., 2021; Reichelt et al., 2021).

This work adds to the already-known role of human and rodent EC➔hippocampal projections in learning and memory (Yassa et al., 2010; Stone et al., 2011; Wilson et al., 2013; Hansen et al., 2018; Yun et al., 2018b; Amani et al., 2021), antidepressant-like behavior (Yun et al., 2018a), and reward-seeking (Ge et al., 2017). This work also raises interesting questions. For example, although our work shows that increased activity of LEC➔DG projections does not change motivation for an operant reward, other work shows a role for midbrain dopamine (a neurotransmitter often linked to salience and reward) in EC associative memory encoding (Lee et al., 2021). Future research is warranted into whether the activity of LEC➔DG projections also regulate behavioral pattern separation and cognitive flexibility in the context of animal states and traits, such as animal models for addiction, which are marked by altered midbrain dopaminergic neuron activity (Schultz, 1997; Ungless et al., 2010; Morikawa and Paladini, 2011; Marinelli and McCutcheon, 2014; Keiflin and Janak, 2015; Wise and Jordan, 2021), and whether different EC➔DG or EC➔hippocampal circuits are engaged under different conditions (Kassab and Alexandre, 2018). Also, while the LEC➔DG projections stimulated in the present work are glutamatergic, GABAergic EC➔hippocampus projections also exist (Melzer et al., 2012; Caputi et al., 2013; Basu et al., 2016). These long-range inhibitory EC➔hippocampus neurons synapse are positioned to disinhibit hippocampal activity as they synapse on hippocampal GABAergic neurons. Thus, it would be interesting to see how modulated activity of long-range inhibitory EC➔hippocampal projections influences behavioral pattern separation and cognitive flexibility.

In sum, the present data show altered activity of mouse LEC➔DG projections regulates both behavioral pattern separation and cognitive flexibility. This work provides insight into the cellular and circuit mechanisms underlying these cognitive abilities and opens avenues for developing circuit-based treatments for impaired hippocampal cognition. As such, these data advance fundamental and translational neuroscience knowledge relevant to two cognitive functions critical for adaptation and survival.

## Data Availability Statement

Raw data will be placed in two archives: MouseBytes (https://www.mousebytes.ca/home) is available on written request.

## Ethics Statement

The animal study was reviewed and approved by the IACUC — Children’s Hospital of Philadelphia.

## Author Contributions (based on Project CRediT)

*initials listed via author order on study*

Conceptualization: SY, AJE, RPR

Methodology: IS, SY, FTH, MS, RPR, AJE

Software: RS, GLB

Validation: SY, HAH, GLB, AJE

Formal Analysis: IS, FTH, SY, HAH,

GLB Investigation: IS, SY, FHT, MS, CdS, RPR, MS, HAH

Resources: SY, AJE

Data Curation: IS, SY, HAH, GLB

Writing, original draft: SY, AJE

Writing, review and editing: SY, AJE, HAH

Visualization: SY, HAH

Supervision: SY, AJE

Projection Administration: SY, AJE

Funding Acquisition: SY, AJE

## Acknowledgements

We thank many scientists for technical support and helpful conversations including Kyung Jin Ahn and Dr. Ann Stowe. We thank staff members of Eisch lab members who help make our experiments possible.

## Data Availability Statement

Raw data will be placed in two archives: MouseBytes (https://www.mousebytes.ca/home) is also available on written request.

## Ethics

Human subjects: No

Animal subjects: Yes

Ethics statement: The study was approved by three Ethics committees (the Institutional Animal Care and Use Committees at the University of Texas Southwestern Medical Center [UTSW], Children’s Hospital of Philadelphia [CHOP], and Brookhaven National Laboratories [BNL]). Specifically, animal procedures and husbandry were in accordance with the National Institutes of Health Guide for the Care and Use of Laboratory Animals, and performed in IACUC-approved facilities at UT Southwestern Medical Center (UTSW, Dallas TX; AAALAC Accreditation #000673, PHS Animal Welfare Assurance D16-00296, Office of Laboratory Animal Welfare [OLAW] A3472-01), Children’s Hospital of Philadelphia (CHOP, Philadelphia, PA; AAALAC Accreditation #000427, PHS Animal Welfare Assurance D16-00280 [OLAW A3442-01]) and Brookhaven National Laboratories (BNL, Upton NY; AAALAC Accreditation #000048, PHS Animal Welfare Assurance D16-00067 [OLAW A3106-01]).

## FIGURE LEGENDS

**Supplementary Figure 1. Relationship between hippocampal terminal regions expressing viral-mediated green fluorescent protein (GFP) and performance in LDR Test. (A-D)** Fourteen weeks after bilateral LEC stereotaxic infusion of AAV-TRIP8bshRNA-EGFP or AAV-SCRshRNA-EGFP, GFP-immunoreactive (GFP+) cell bodies were detected in LEC Layer IIa **(A)**, a layer enriched with stellate cells that project to the dentate gyrus molecular layer, DG Mol) and IIb **(C)**, a layer enriched with pyramidal cells that project to the CA1 stratum lacunosum molecular [SLM]). In mice in which the LEC IIa was targeted, GFP+ terminals were detected in the outer DG Mol **(B)**; such mice were termed LEC=>DG Mol mice. If the virus also spread to LEC IIb, then GFP+ terminals were evident in both the DG Mol as well as the CA1 SLM **(D)**; these mice were termed LEC➔DG Mol+CA1 SLM mice. Data in main text figures are all from LEC➔DG Mol mice (mice with GFP+ terminal expression in the DG Mol but no expression in the CA1 SLM). In **(E-J)**, behavioral data is presented showing SCR shRNA mice and TRIP8b shRNA mice from both LEC➔DG Mol mice and LEC➔DG Mol+CA1 SLM mice. In Large separation, SCR shRNA and TRIP8b shRNA mice (both LEC➔DG Mol and LEC➔DG Mol+CA1 SLM) had similar LDR Test measures, including **(E)** time to 1st reversal, **(F)** % correct to 1st reversal, and **(G)** reversal #. However, in Small separation, TRIP8b shRNA LEC=>DG Mol mice had better pattern separation vs. SCR shRNA based on **(H)** time to 1st reversal and **(I)** % correct to 1st reversal but performed similarly in reversal learning **(J)**. Dotted lines delineate **(A, C)** IIa and IIb and **(B, D)** DG granule cell layer (GCL). Scale bar **(A)**=200 um applies **(A-D)**. One-way ANOVA used for all. **(E)** Main Effect: Treatment F(2, 21)=0.3079, p=0.7383. **(F)** Main Effect: Treatment F(2, 21)=0.4144, p=0.6660. **(G)** Main Effect: Treatment F(2, 21)=0.8267, p=0.4512. **(H)** Main Effect: Treatment F(2, 21)=6.406, **p=0.0067, *post hoc*: **p=0.0068 in SCR vs TRIP8b mice (LEC➔DG Mol mice). **(I)** Main Effect: Treatment F(2, 21)=3.496, *p=0.0489, *post hoc*: *p=0.0472 in SCR vs. TRIP8b mice (LEC➔DG Mol mice). **(J)** Main Effect: Treatment F(2, 21)=2.882, p=0.0783. Complete statistical information provided in **Supplementary Table 1**.

