## Supplementary table 1 for "Behavioral pattern separation and cognitive flexibility are enhanced in a mouse model of increased lateral entorhinal cortex-dentate gyrus circuit activity"

**Supplementary Table 1. Statistical analyses for each figure panel and all results.**

Bold text and ***, p<0.05. Italicized text, 0.05<p<0.1. *Magnitudes of partial omega-squared (for RM two-way ANOVA): 0.01 small; 0.06 medium; 0.14 large. N/A not applicable.

| Dependent Measure | Figure Panel | Group: Subject Number | Mean | | | | | | | | | | | | | | | | Statistical Test | Variables, Main Effect, Interaction *P<0.5 **P<0.01 ***P<0.001 ****P<0.0001 | F Value | P value | Post hoc Test (Bonferroni for 2-way ANOVA, Tukey for 1-way ANOVA) | Effect size (when RM two-way ANOVA p<0.05, partial omega-squared is calculated where ≤0.05 small, ≥0.06 medium, ≥0.14 large) | CI (95%) |
|---|---|---|---|---|---|---|---|---|---|---|---|---|---|---|---|---|---|---|---|---|---|---|---|---|---|
| Weights | 1D | | Post-surgery | 0 | 1 | 2 | 3 | 4 | 5 | 6 | 7 | 8 | 9 | 10 | 11 | 12 | 13 | 14 | | | | | | | |
| | | SCR: 9 | SCR | 25.2 | 25.77 | 24.71 | 24.98 | 25.46 | 25.9 | 26.28 | 26.36 | 26.76 | 26.96 | 27.21 | 27.14 | 27.51 | 28.04 | 32.04 | RM 2-way ANOVA | Time**** | F (14, 238) = 130.6 | **P<0.0001** | N/A | | |
| | | TRIP8b: 10 | TRIP | 24.43 | 25.98 | 24.27 | 24.54 | 24.62 | 24.95 | 25.45 | 25.5 | 26.21 | 26.21 | 26.78 | 26.93 | 27.1 | 27.65 | 31.41 | | Virus | F (1, 17) = 0.6704 | P=0.4242 | | | |
| | | | | | | | | | | | | | | | | | | | | Interaction | F (14, 238) = 0.9406 | P=0.5158 | | | |
| General Touchscreen Training w/six windows | 2A | SCR: 9 | General TS training | | | | | | | | | | | | | | | | | Training | | | | | |
| | | | Training Stage | | HAB | | | IT | | | MT | | | MI | | | PI | | RM 2-way ANOVA | Stage**** | F (4, 68) = 75.03 | **P<0.0001** | N/A | 0.92 | .77, .96 |
| | | TRIP8b: 10 | SCR | | 1 | | | 1 | | | 5.222 | | | 1.111 | | | 12.22 | | | Virus | F (1, 17) = 0.6292 | P=0.4386 | | | |
| | | | TRIP | | 1 | | | 1 | | | 4.4 | | | 1.1 | | | 10.9 | | | Interaction | F (4, 68) = 0.3333 | P=0.8547 | | | |
| Punish Incorrect: Session Length | 2B | SCR: 9 | Session length (s) | | | | | | | | | | | | | | | | | Training | | | | | |
| | | | Session | | | | Day 1 | | | | | | Last | | | | | | RM 2-way ANOVA | Stage | F (1, 17) = 0.0857 | P=0.7731 | N/A | | |
| | | TRIP8b: 10 | SCR | | | | 1292 | | | | | | 1256 | | | | | | | Virus | F (1, 17) = 0.2492 | P=0.6240 | | | |
| | | | TRIP | | | | 1349 | | | | | | 1312 | | | | | | | Interaction | F (1, 17) = 2.128e | P=0.9964 | | | |
| Punish Incorrect: Trial Number | 2C | SCR: 9 | Trial # | | | | | | | | | | | | | | | | | Training | | | | | |
| | | | Session | | | | Day 1 | | | | | | Last | | | | | | RM 2-way ANOVA | Stage | F (1, 17) = 2.858 | P=0.1092 | N/A | | |
| | | TRIP8b: 10 | SCR | | | | 29.89 | | | | | | 30 | | | | | | | Virus | F (1, 17) = 2.335 | P=0.1449 | | | |
| | | | TRIP | | | | 27.8 | | | | | | 30 | | | | | | | Interaction | F (1, 17) = 2.335 | P=0.1449 | | | |
| Punish Incorrect: Percent Correct | 2D | SCR: 9 | % correct | | | | | | | | | | | | | | | | | Training | | | | | |
| | | | Session | | | | Day 1 | | | | | | Last | | | | | | RM 2-way ANOVA | Stage**** | F (1, 17) = 148.2 | **P<0.0001** | Day 1 vs Last on both: ****p<0.0001 | 0.85 | .64, .94 |
| | | TRIP8b: 10 | SCR | | | | 53.19 | | | | | | 90 | | | | | | | Virus | F (1, 17) = 1.599 | P=0.2231 | | | |
| | | | TRIP | | | | 61.82 | | | | | | 89 | | | | | | | Interaction | F (1, 17) = 3.358 | P=0.0844 | | | |
| LDR Train: Percent reaching criteria | 3B | | LD train criteria completion curve | | | | | | | | | | | | | | | | Log-rank (Mantel-Cox) test | N/A | N/A | p=0.5328 | N/A | | |
| | | | SCR | | | | | | | | Median: 5.5 | | | | | | | | | | | | | | |
| | | | TRIP | | | | | | | | Median: 7 | | | | | | | | | | | | | | |
| LDR Train: Days to completion | 3C | | Days to completion | | | | | | | | | | | | | | | | Unpaired T-test | N/A | N/A | p=0.1088 | N/A | | |
| | | SCR: 9 | SCR | | | | | | | | 5.111 ± 0.824 | | | | | | | | | | | | | | |
| | | TRIP8b: 10 | TRIP | | | | | | | | 7.4 ± 1.046 | | | | | | | | | | | | | | |
| LDR Train: Percent correct | 3D | SCR: 9 | % correct | | | | | | | | | | | | | | | | | | | | | | |
| | | | Session | | | | Day 1 | | | | | | Last day | | | | | | RM 2-way ANOVA | Day** | F (1, 17) = 10.08 | **p=0.0055** | Day 1 vs Last day in TRIP8b: **p=0.0075 | 0.19 | 0.00, 0.53 |
| | | TRIP8b: 10 | SCR | | | | 39.29 | | | | | | 45.47 | | | | | | | Virus | F (1, 17) = 0.0006 | p=0.9796 | | | |
| | | | TRIP | | | | 34.87 | | | | | | 50.42 | | | | | | | Interaction | F (1, 17) = 2.084 | p=0.1671 | | | |
| LDR Train: Time to 1st reversal | 3E | SCR: 9 | Time to 1st reversal | | | | | | | | | | | | | | | | | | | | | | |
| | | | Session | | | | Day 1 | | | | | | Last day | | | | | | RM 2-way ANOVA | Day | F (1, 17) = 1.386 | p=0.2552 | N/A | | |
| | | TRIP8b: 10 | SCR | | | | 1316 | | | | | | 760 | | | | | | | Virus | F (1, 17) = 1.500 | p=0.2373 | | | |
| | | | TRIP | | | | 857.6 | | | | | | 932.3 | | | | | | | Interaction | F (1, 17) = 2.382 | p=0.1411 | | | |
| LDR Train: Percent correct to 1st reversal | 3F | SCR: 9 | % correct to 1st reversal | | | | | | | | | | | | | | | | | | | | | | |
| | | | Session | | | | Day 1 | | | | | | Last day | | | | | | RM 2-way ANOVA | Day | F (1, 17) = 1.571 | p=0.2271 | N/A | | |
| | | TRIP8b: 10 | SCR | | | | 47.92 | | | | | | 64.97 | | | | | | | Virus | F (1, 17) = 2.700 | p=0.1187 | | | |
| | | | TRIP | | | | 63.57 | | | | | | 68.08 | | | | | | | Interaction | F (1, 17) = 0.5309 | p=0.4762 | | | |
| LDR Test: Block 1, time to 1st reversal | 4C | SCR: 9 | Block 1 time to 1st reversal (sec) | | | | | | | | | | | | | | | | | | | | | | |
| | | | Session | | | | Large | | | | | | Small | | | | | | RM 2-way ANOVA | Separation | F (1, 17) = 1.084 | p=0.3124 | N/A | | |
| | | TRIP8b: 10 | SCR | | | | 1032 | | | | | | 1460 | | | | | | | Virus | F (1, 17) = 2.423 | p=0.1380 | | | |
| | | | TRIP | | | | 1296 | | | | | | 1380 | | | | | | | Interaction | F (1, 17) = 0.1799 | p=0.6768 | | | |
| LDR Test: Block 1, percent correct to 1st reversal | 4D | SCR: 9 | Block 1 % correct to 1st reversal | | | | | | | | | | | | | | | | | Separation | | | | | |
| | | | Session | | | | Large | | | | | | Small | | | | | | RM 2-way ANOVA | | F (1, 17) = 1.679 | p=0.2123 | N/A | | |
| | | TRIP8b: 10 | SCR | | | | 61.69 | | | | | | 47.32 | | | | | | | Virus | F (1, 17) = 0.3233 | p=0.5771 | | | |
| | | | TRIP | | | | 52.42 | | | | | | 50.03 | | | | | | | Interaction | F (1, 17) = 0.8583 | p=0.3672 | | | |
| LDR Test: Block 6, time to 1st reversal | 4E | SCR: 9 | Block 6 time to 1st reversal (sec) | | | | | | | | | | | | | | | | | Separation*** | | | | | |
| | | | Session | | | | Large | | | | | | Small | | | | | | RM 2-way ANOVA | | F (1, 17) = 16.36 | **p=0.0008** | SCR vs TRIP8b in Small Separation: **p=0.0012 | 0.37 | 0.04, 0.68 |
| | | TRIP8b: 10 | SCR | | | | 730.6 | | | | | | 1685 | | | | | | | Virus** | F (1, 17) = 15.53 | **p=0.0011** | | 0.19 | 0.00, 0.53 |
| | | | TRIP | | | | 604.8 | | | | | | 950.5 | | | | | | | Interaction | F (1, 17) = 3.589 | p=0.0753 | | | |
| LDR Test: Block 6, percent correct to 1st reversal | 4F | SCR: 9 | Block 6 % correct to 1st reversal | | | | | | | | | | | | | | | | | Separation | | | | | |
| | | | Session | | | | Large | | | | | | Small | | | | | | RM 2-way ANOVA | | F (1, 17) = 1.760 | p=0.2022 | SCR vs TRIP8b in Small Separation: *p=0.0397 | | |
| | | TRIP8b: 10 | SCR | | | | 59.29 | | | | | | 44.03 | | | | | | | Virus** | F (1, 17) = 8.939 | **p=0.0082** | | 0.10 | 0.00, 0.44 |
| | | | TRIP | | | | 66.31 | | | | | | 63.76 | | | | | | | Interaction | F (1, 17) = 0.8945 | p=0.3575 | | | |
| LDR Test: Block 1, reversal # | 4G | SCR: 9 | Block 1 reversal # | | | | | | | | | | | | | | | | | Separation** | | | | | |
| | | | Session | | | | Large | | | | | | Small | | | | | | RM 2-way ANOVA | | F (1, 17) = 10.07 | **p=0.0056** | Large Separation vs Small Separation in SCR: *p=0.0156 | 0.12 | 0.00, 0.46 |
| | | TRIP8b: 10 | SCR | | | | 1.444 | | | | | | 0.5556 | | | | | | | Virus | F (1, 17) = 1.448 | p=0.2454 | | | |
| | | | TRIP | | | | 0.8 | | | | | | 0.4 | | | | | | | Interaction | F (1, 17) = 1.448 | p=0.2453 | | | |
| LDR Test: Block 6, reversal # | 4H | SCR: 9 | Block 6 reversal # | | | | | | | | | | | | | | | | | Separation** | | | | | |
| | | | Session | | | | Large | | | | | | Small | | | | | | RM 2-way ANOVA | | F (1, 17) = 14.83 | **p=0.0013** | SCR vs TRIP8b in Small separation: #p=0.0814 | 0.26 | 0.00, 0.59 |
| | | TRIP8b: 10 | SCR | | | | 1.556 | | | | | | 0.3333 | | | | | | | Virus* | F (1, 17) = 5.531 | **p=0.0310** | | 0.12 | 0.00, 0.46 |
| | | | TRIP | | | | 2.1 | | | | | | 1.2 | | | | | | | Interaction | F (1, 17) = 0.3419 | p=0.5664 | | | |
| LDR Test: Block 1, correct image choice latency | 5A | SCR: 9 | Block 1 correct image choice latency (sec) | | | | | | | | | | | | | | | | | | | | | | |
| | | | Session | | | | Large | | | | | | Small | | | | | | RM 2-way ANOVA | Separation | F (1, 17) = 2.977 | p=0.1026 | SCR vs TRIP8b in Small separation: p=0.1204 | | |
| | | TRIP8b: 10 | SCR | | | | 9.151 | | | | | | 7.497 | | | | | | | Virus | F (1, 17) = 0.4937 | p=0.4918 | | | |
| | | | TRIP | | | | 3.87 | | | | | | 19.06 | | | | | | | Interaction* | F (1, 17) = 4.610 | **p=0.0465** | | 0.08 | 0.00, 0.41 |
| LDR Test: Block 6, correct image choice latency | 5B | SCR: 9 | Block 6 correct image choice latency (sec) | | | | | | | | | | | | | | | | | | | | | | |
| | | | Session | | | | Large | | | | | | Small | | | | | | RM 2-way ANOVA | Separation | F (1, 17) = 0.0003 | p=0.9855 | N/A | | |
| | | TRIP8b: 10 | SCR | | | | 13.24 | | | | | | 8.5 | | | | | | | Virus | F (1, 17) = 0.8853 | p=0.3599 | | | |
| | | | TRIP | | | | 3.68 | | | | | | 8.619 | | | | | | | Interaction | F (1, 17) = 0.8399 | p=0.3722 | | | |
| LDR Test: Block 1, reward collection latency | 5C | SCR: 9 | Block 1 reward collection latency (sec) | | | | | | | | | | | | | | | | | | | | | | |
| | | | Session | | | | Large | | | | | | Small | | | | | | RM 2-way ANOVA | Separation | F (1, 17) = 1.144 | p=0.2997 | N/A | | |
| | | TRIP8b: 10 | SCR | | | | 1.376 | | | | | | 2.89 | | | | | | | Virus | F (1, 17) = 2.248 | p=0.1521 | | | |
| | | | TRIP | | | | 1.217 | | | | | | 1.105 | | | | | | | Interaction | F (1, 17) = 1.539 | p=0.2316 | | | |
| LDR Test: Block 6, reward collection latency | 5D | SCR: 9 | Block 6 reward collection latency (sec) | | | | | | | | | | | | | | | | | | | | | | |
| | | | Session | | | | Large | | | | | | Small | | | | | | RM 2-way ANOVA | Separation | F (1, 17) = 0.0901 | p=0.7676 | N/A | | |
| | | TRIP8b: 10 | SCR | | | | 1.29 | | | | | | 1.632 | | | | | | | Virus | F (1, 17) = 0.1240 | p=0.7291 | | | |
| | | | TRIP | | | | 1.484 | | | | | | 1.281 | | | | | | | Interaction | F (1, 17) = 1.383 | P=0.2559 | | | |
| LDR Test: Block 1, total ITI blank touches | 5E | SCR: 9 | Block 1 total ITI blank touches | | | | | | | | | | | | | | | | | | | | | | |
| | | | Session | | | | Large | | | | | | Small | | | | | | RM 2-way ANOVA | Separation | F (1, 17) = 3.077 | p=0.0974 | SCR vs TRIP8b in Large separation: p=0.1537 | | |
| | | TRIP8b: 10 | SCR | | | | 11.33 | | | | | | 12.67 | | | | | | | Virus | F (1, 17) = 0.7417 | p=0.4011 | | 0.05 | 0.00, 0.36 |
| | | | TRIP | | | | 24 | | | | | | 10.3 | | | | | | | Interaction* | F (1, 17) = 4.547 | **p=0.0478** | | | |
| LDR Test: Block 6, total ITI blank touches | 5F | SCR: 9 | Block 6 total ITI blank touches | | | | | | | | | | | | | | | | | | | | | | |
| | | | Session | | | | Large | | | | | | Small | | | | | | RM 2-way ANOVA | Separation | F (1, 17) = 1.877 | p=0.1885 | N/A | | |
| | | TRIP8b: 10 | SCR | | | | 17.11 | | | | | | 13.67 | | | | | | | Virus | F (1, 17) = 0.0887 | p=0.7694 | | | |
| | | | TRIP | | | | 16 | | | | | | 11.3 | | | | | | | Interaction | F (1, 17) = 0.0446 | p=0.8353 | | | |
| EPM: Total distance traveled | 6A | | | | | | | | Total distance traveled | | | | | | | | | | | | | | | | |
| | | SCR:9 | | | | | | | | 1406 | | | | | | | | | Unpaired t-test | N/A | N/A | p=0.6529 | N/A | | |
| | | TRIP8b: 8 | | | | | | | | 1347 | | | | | | | | | | | | | | | |
| EPM: Duration in | | | Duration in open arms | | | | | | | | | | | | | | | | | | | | | | |

| | | | | | | | | | | | | | | | | |
|---|---|---|---|---|---|---|---|---|---|---|---|---|---|---|---|---|
| EPM: Duration in open arms | 6B | SCR:9 | | | | 90.46 | | | | Unpaired t-test | N/A | N/A | p=0.5044 | N/A | | |
| | | TRIP8b: 8 | | | | 82.7 | | | | | | | | | | |
| EPM: Frequency in open arms | 6C | | | | | Frequency in open arms | | | | | | | | | | |
| | | SCR:9 | | | | 11.33 | | | | Unpaired t-test | N/A | N/A | p=0.9863 | N/A | | |
| | | TRIP8b: 8 | | | | 11.38 | | | | | | | | | | |
| Total DCX+ cells | 7C | | | | | | | | | 2-way Anova | | | | | | |
| | | | All cells | | Progenitors | | | | Immature Cells | | Cell type**** | F (2, 45) = 165.4 | **P<0.0001** | | | |
| | | SCR:9 | 5685 | | 2260 | | | | 3426 | | Virus**** | F (1, 45) = 23.58 | **P<0.0001** | SCR vs TRIP8b in Immature Cells: p=0.0292 | 0.57 | 0.35, 0.73 |
| | | TRIP8b: 8 | 6936 | | 2846 | | | | 4098 | | Intreraction | F (2, 45) = 1.466 | P=0.2416 | | | |
| DCX+ cells (superior vs inferior) | 7D | | | Superior | | | | Inferior | | 2-way Anova | Regions* | F (1, 30) = 6.466 | **P=0.0164** | SCR vs TRIP8B in superior region: p=0.0374 | | |
| | | SCR:9 | | 3113 | | | | 2572 | | | Virus*** | F (1, 30) = 14.50 | **P=0.0006** | SCR vs TRIP8B in inferior region: p=0.0032 | 0.24 | 0.03, 0.51 |
| | | TRIP8b: 8 | | 3622 | | | | 3322 | | | Intreraction | F (1, 30) = 0.5293 | P=0.4726 | | | |
| DCX+ progenitor cells (superior vs inferior) | 7E | | | Superior | | | | Inferior | | 2-way Anova | Regions | F (1, 30) = 1.712 | P=0.2006 | | | |
| | | SCR:9 | | 1241 | | | | 1019 | | | Virus** | F (1, 30) = 10.55 | **P=0.0029** | SCR vs TRIP8b in inferior region: p=0.0041 | 0.36 | 0.10, 0.61 |
| | | TRIP8b: 8 | | 1430 | | | | 1416 | | | Intreraction | F (1, 30) = 1.330 | P=0.2579 | | | |
| DCX+ immature cells (superior vs inferior) | 7F | | | Superior | | | | Inferior | | 2-way Anova | Regions* | F (1, 30) = 7.121 | **P=0.0122** | SCR vs TRIP8b in superior region: p=0.0548 | 0.31 | 0.07, 0.57 |
| | | SCR:9 | | 1872 | | | | 1554 | | | Virus** | F (1, 30) = 8.813 | **P=0.0058** | SCR vs TRIP8b in inferior region: p=0.0357 | | |
| | | TRIP8b: 8 | | 2192 | | | | 1906 | | | Intreraction | F (1, 30) = 0.0202 | P=0.8878 | | | |
| LDR Test: Block 6, Large separation, time to 1st reversal | Supp Fig 1E | | Block 6 Large separation Time to 1st reversal (sec) | | | | | | | 1-way ANOVA | Treatment | F (2, 21) = 0.3079 | P=0.7383 | N/A | | |
| | | SCR: 9 | SCR | | | | 730.6 | | | | | | | | | |
| | | TRIP8b (Mol): 10 | TRIP 8b (Mol) | | | | 604.8 | | | | | | | | | |
| | | TRIP8b (Mol + CA1): 10 | TRIP 8b (Mol + CA1) | | | | 575.8 | | | | | | | | | |
| LDR Test: Block 6, Large separation, percent correct to 1st reversal | Supp Fig 1F | | Block 6 Large separation % correct to 1st reversal | | | | | | | 1-way ANOVA | Treatment | F (2, 21) = 0.4144 | P=0.6660 | N/A | | |
| | | SCR: 9 | SCR | | | | 59.29 | | | | | | | | | |
| | | TRIP8b (Mol): 10 | TRIP 8b (Mol) | | | | 66.31 | | | | | | | | | |
| | | TRIP8b (Mol + CA1): 10 | TRIP 8b (Mol + CA1) | | | | 60.91 | | | | | | | | | |
| LDR Test: Block 6, Large separation, reversal number | Supp Fig 1G | | Block 6 Large separation reversal # | | | | | | | 1-way ANOVA | Treatment | F (2, 21) = 0.8267 | P=0.4512 | N/A | | |
| | | SCR: 9 | SCR | | | | 1.556 | | | | | | | | | |
| | | TRIP8b (Mol): 10 | TRIP 8b (Mol) | | | | 2.1 | | | | | | | | | |
| | | TRIP8b (Mol + CA1): 10 | TRIP 8b (Mol + CA1) | | | | 1.8 | | | | | | | | | |
| LDR Test: Block 6, Small separation, time to 1st reversal | Supp Fig 1H | | Block 6 Small separation Time to 1st reversal (sec) | | | | | | | 1-way ANOVA | Treatment** | F (2, 21) = 6.406 | **P=0.0067** | SCR vs TRIP8b (Mol): **p=0.0068 | 0.31 | 0.01, 0.58 |
| | | SCR: 9 | SCR | | | | 1685 | | | | | | | | | |
| | | TRIP8b (Mol): 10 | TRIP 8b (Mol) | | | | 950.5 | | | | | | | | | |
| | | TRIP8b (Mol + CA1): 10 | TRIP 8b (Mol + CA1) | | | | 1060 | | | | | | | | | |
| LDR Test: Block 6, Small separation, percent correct to 1st reversal | Supp Fig 1I | | Block 6 Small separation % correct to 1st reversal | | | | | | | 1-way ANOVA | Treatment* | F (2, 21) = 3.496 | **P=0.0489** | SCR vs TRIP8b (Mol): *p=0.0472 | 0.17 | 0.00, 0.45 |
| | | SCR: 9 | SCR | | | | 44.03 | | | | | | | | | |
| | | TRIP8b (Mol): 10 | TRIP 8b (Mol) | | | | 63.76 | | | | | | | | | |
| | | TRIP8b (Mol + CA1): 5 | TRIP 8b (Mol + CA1) | | | | 48.46 | | | | | | | | | |
| LDR Test: Block 6, Small separation, reversal number | Supp Fig 1J | | Block 6 Small separation reversal # | | | | | | | 1-way ANOVA | Treatment | F (2, 21) = 2.882 | P=0.0783 | N/A | | |
| | | SCR: 9 | SCR | | | | 0.3333 | | | | | | | | | |
| | | TRIP8b (Mol): 10 | TRIP 8b (Mol) | | | | 1.2 | | | | | | | | | |
| | | TRIP8b (Mol + CA1): 5 | TRIP 8b (Mol + CA1) | | | | 1 | | | | | | | | | |
