## Supplementary figures and images for "Behavioral pattern separation and cognitive flexibility are enhanced in a mouse model of increased lateral entorhinal cortex-dentate gyrus circuit activity"

### Supplementary figure 1

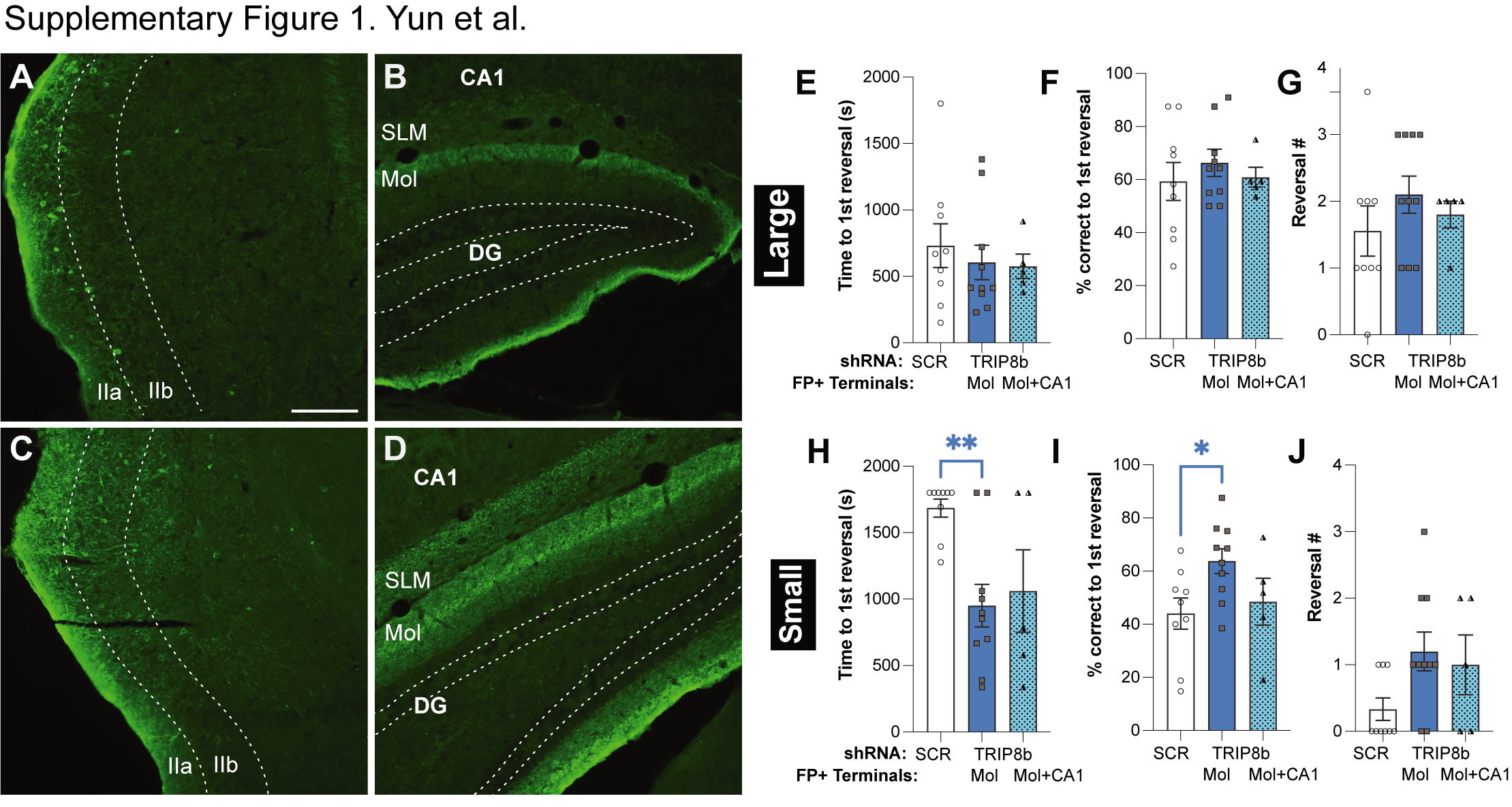
